# A single dose of the antipsychotic drug clozapine has long-term behavioral and functional effects in mice

**DOI:** 10.64898/2026.03.27.714783

**Authors:** Leonardo Lupori, Matthias Heindorf, Stylianos Kouvaros, Tristan Schilling, Alice Schildkamp, Josef Bischofberger, Georg B. Keller

## Abstract

Antipsychotic dosing regimens, commonly daily or via slow-release compounds, are designed to maintain steady-state plasma concentrations. They are guided by their plasma half-life, which is typically in the range of several hours. This schedule contrasts with the slow time course of therapeutic efficacy, which takes days to weeks to develop fully. This discrepancy led us to hypothesize that the effects of a single dose of an antipsychotic drug might be detectable well beyond the time window predicted by receptor occupancy. To test this, we administered a single dose of the antipsychotic drug clozapine to mice. We observed long-term behavioral effects and changes to cortical activity patterns up to several days after administration. Specifically, clozapine induced a decorrelation of activity in the dorsal cortex observable up to 9 days post administration. This effect was driven by a genetically and functionally distinct subset of layer 5 intratelencephalic neurons and was accompanied by an increase in the reliability of long-range inhibitory functional influence on a similar timescale. Thus, a single dose of clozapine triggers changes in the function of specific circuits that outlast the presence of the drug. If antipsychotic action depends on slower processes involving plasticity, longer dosing intervals may warrant further clinical exploration.

## INTRODUCTION

Schizophrenia is typically treated with a class of medications known as antipsychotic (AP) drugs. Despite decades of clinical use, how these drugs change the function of neuronal circuits, and which of these changes underlie their therapeutic effect, remains unclear. By contrast, drug-receptor interactions have been extensively characterized both in vitro (Roth et al. 2004; McCutcheon et al. 2023) and in patients (Kapur et al. 2000; Hart et al. 2024), and they guide how AP drugs are used in the clinic.

Dosing regimens are typically determined by correlating plasma concentration with clinical efficacy and receptor occupancy (Byerly and DeVane 1996; Northwood et al. 2023; Hart et al. 2024). This approach assumes that therapeutic effect follows the presence of the drug at its target receptors. However, several lines of evidence challenge this assumption. For AP drugs, receptor occupancy reaches levels comparable to those of continuous treatment within hours of the first dose (Nordström et al. 1992; Nyberg et al. 1997). Clinical improvement, by contrast, starts within the first week of treatment and continues to build up over the following weeks (Agid et al. 2003, 2006), long after receptor occupancy has stabilized.

The slow time course also shapes clinical practice: Before concluding that a patient does not respond to an AP drug, clinicians have to give it at an adequate dose for at least six weeks (Howes et al. 2017). Patients whose symptoms persist despite at least two such treatment attempts are diagnosed with treatment-resistant schizophrenia (Howes et al. 2017), which affects up to one third of patients (Demjaha et al. 2017; Siskind et al. 2022). For these patients, clozapine is the only recommended drug (Keepers et al. 2020; Kane et al. 1988), and it is one of the most effective AP drugs overall (Efthimiou et al. 2024). Yet, even with clozapine, clinical improvement continues to build up over several weeks and, in some patients, for up to one year after the start of treatment (Meltzer et al. 1989, 1990; Meltzer 1992; Wilson 1996).

One explanation for why clinical improvement develops so much more slowly than receptor occupancy is that the drug’s action at its targets is only the first step of a slower process, likely involving neuronal plasticity, that gradually reshapes circuit function (Konradi and Heckers 2001). This hypothesis predicts that some treatment-relevant effects of AP drugs on neuronal circuits should outlast the presence of the drug in the brain. We expected that such effects would involve long-range communication in cortex, which is central to recent computational accounts of psychosis (Sterzer et al. 2018; Keller and Sterzer 2024).

To test this prediction, we used a single dose of clozapine in mice. In rodents, clozapine is cleared from plasma and brain within a few hours (Baldessarini et al. 1993; Jendryka et al. 2019), allowing us to isolate long-term effects that the drug initiates. We characterized alterations in behavior as well as changes in cortical activity patterns and functional influence over the course of several days after drug administration. We show that, beyond its acute effects, a single dose of clozapine is sufficient to modulate behavior and cortical dynamics for up to 9 days. These changes were accompanied by alterations in cortico-cortical and thalamo-cortical functional connectivity on a similar timescale, consistent with a plasticity-driven reshaping of circuit function. Finally, we show that long-term functional effects of a single dose generalize to three additional AP drugs with different pharmacological profiles (aripiprazole, risperidone, and haloperidol). Because these effects do not depend on the continued presence of the drug, we speculate that they are candidates for the circuit changes underlying AP action.

## RESULTS

### A single dose of clozapine induces long-term behavioral changes

We first asked whether we could find evidence of behavioral changes in mice following a single dose of clozapine. To do this, we performed an open field test and measured features of spontaneous behavior before and at different time points after a single clozapine injection. We compared three experimental groups of mice that received either a saline injection, a low dose of 1 mg/kg, or a standard dose of 10 mg/kg (**Figure 1A**). As mice explored the arena, we tracked the position of a set of body parts (**Figure 1A**) (Mathis et al. 2018). We first quantified exploratory behavior by analyzing the trajectory of the nose across sessions (**Figure 1B**). Consistent with previous reports, we found that clozapine treatment acutely induced a sharp reduction in total distance traveled one hour after injection in a dose-dependent manner. This acute sedative effect of clozapine is well described in rodents (Ballard and Mcallister 1999; McOmish et al. 2012), and is also observed in patients (Hopkins and Miller 2025; Ramos Perdigués et al. 2016). By 24 hours post administration, the low dose group was no longer different from saline controls while the standard dose group still showed a significant reduction in distance traveled. Surprisingly, however, we found that a week after injection, mice that received 10 mg/kg clozapine traveled a significantly shorter distance than saline controls (**Figure 1C**). To test whether such long-term differences were also apparent in more detailed behavioral patterns, we investigated how spontaneous behavior changed following the injection of clozapine. We segmented each open field session into different behavioral syllables in an unsupervised manner (**Figure 1D**) (Weinreb et al. 2024). We measured two aspects of behavioral structure: the frequency of each syllable, and the transition probability between all syllable pairs organized into a transition matrix. To compare treatment groups, we calculated the L1 distance between either the mean syllable frequency vectors or the mean transition matrices of each group (**Methods**) and compared these values with a null distribution of distances obtained by randomly shuffling the treatment labels (**Figure 1E-H**, bottom) (von Ziegler et al. 2024). To qualitatively visualize the dataset, we projected the data of each mouse in a two-dimensional space by reducing the dimensionality of either syllable frequencies or transitions using t-distributed stochastic neighbor embedding (t-SNE) (**Figure 1E-H**, top). A low dose of clozapine was sufficient to alter both the frequency and the transition probability of behavioral components one hour after the injection as the distance between treatment groups was higher than expected by chance (**Figure 1E, G**). A higher dose induced robust alterations in both metrics, not only acutely, but also up to 9 days after injection (**Figure 1F, H**). This time course was similar to that observed in the clozapine-driven reduction in total distance traveled. Additionally, we observed no evidence of an effect of clozapine on anxiety levels as measured by thigmotaxis (**Figure S1**). Taken together, these data show that a single injection of clozapine can give rise to long-term changes in mouse spontaneous behavior.

**Figure 1.**
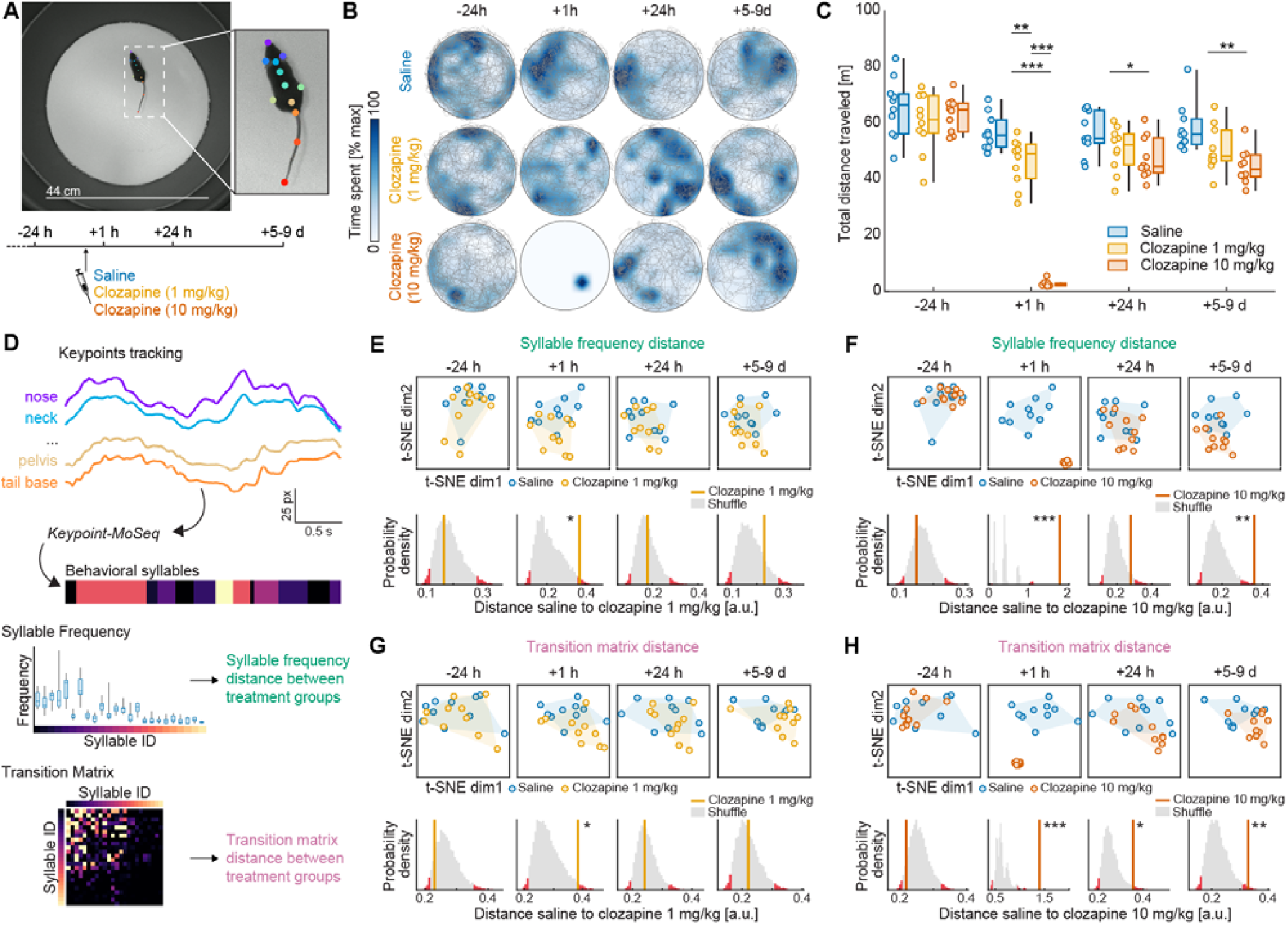
A single dose of clozapine induces long-term behavioral changes. **(A)**Top: Example frame of the open field video recordings with estimated positions of body parts overlaid. Bottom: Time course of the experiment. **(B)**Examples of the trajectories of three mice (nose) at different time points before and after injection of saline or clozapine. **(C)**Total distance traveled per session. Whiskers represent the data range, boxes the interquartile range, and the horizontal line the median. Each data point is one mouse. Here and elsewhere, *: p < 0.05, **: p < 0.01, ***: p < 0.001. See **Table S1** for all statistical information. **(D)**Body part tracking data were used to segment behavior into syllables. For each mouse, we calculated syllable usage frequencies and the frequency of transition between each pair of syllables. The effect of the treatments was quantified as the L1 distance between the mean syllable frequency vectors or the mean transition matrices of saline and clozapine injected mice. **(E)**Top: Saline or clozapine injected mice (1 mg/kg) plotted in a t-SNE embedding space derived from syllable frequency vectors. Shading represents the convex hull of all the points in each group. Bottom: Distribution of the distances for shuffled data. The red bars represent bins below the 2.5 or above the 97.5 percentiles of the shuffle distribution. The orange vertical line represents the observed distance between mice that received saline and clozapine injected (1 mg/kg) mice. **(F)** As in **E**, but for mice that received saline or 10 mg/kg clozapine. **(G)** As in **E**, but using transition matrix data. **(H)** As in **F**, but using transition matrix data.

### A single dose of clozapine induces long-term functional decorrelation in a subset of layer 5 IT neurons

Clozapine administration was shown to decrease the correlation of neuronal activity across dorsal cortex preferentially in a subpopulation of layer 5 (L5) intratelencephalic (IT) neurons (Heindorf and Keller 2024) labeled using the Tlx3-Cre driver line. To test if clozapine also has long-term effects on cortical activity patterns, we performed widefield calcium imaging of L5 IT neurons in dorsal cortex of awake head-fixed mice (**Figure 2A**). We expressed the calcium indicator GCaMP in these neurons by crossing Tlx3-Cre (Gerfen et al. 2013) with Ai148 mice. We parcellated imaging data of dorsal cortex activity into 818 regions of interest (ROIs) and calculated the correlation coefficients of calcium activity for all pairs of ROIs in naive mice and in different intervals up to 9 days after clozapine injection (**Figure 2B, C**). On the day of injection, after recovery from acute sedation, mice showed lower correlation, and this effect was gone 24 hours later. Interestingly, between 4 and 9 days after the injection, mice showed a significant decorrelation of activity in the L5 IT network. This functional phenotype follows the time course of behavioral effects (**Figure 1**) and suggests that a single dose of clozapine induces plasticity that results in long-term changes to both cortical activity patterns and behavior.

**Figure 2.**
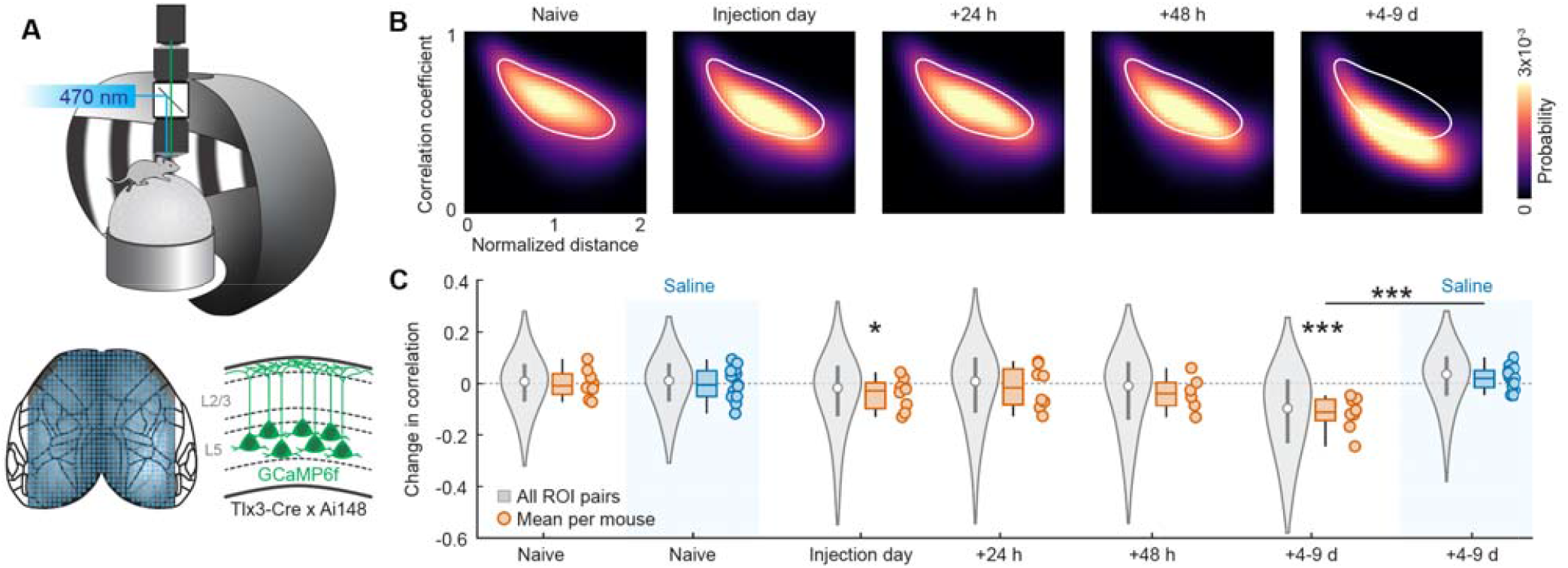
A single dose of clozapine induces long-term decorrelation in the activity of a subset of L5 IT neurons. **(A)**Top: Schematic of the experimental setup. We imaged GCaMP fluorescence using a macroscope. Mice were head-fixed and free to locomote on an air-supported spherical treadmill. Visual stimuli were projected onto a toroidal screen. Bottom left: Dorsal cortex was parcellated with ROIs covering the entire craniotomy. Bottom right: GCaMP6f was expressed in a subset of L5 IT neurons by crossing Tlx3-Cre with Ai148 mice. **(B)** Probability distribution heatmaps of correlation coefficients for all pairs of ROIs as a function of normalized (lambda-bregma) distance between them. The white contour line represents 50% of the maximum probability of the naive time point. **(C)** Time course of the change in correlation with respect to baseline in clozapine injected mice. Change in correlation in saline treated mice is shown for the naive and +4-9 days for comparison. Violin plots represent the distribution of correlation changes for all ROIs (from 1 to 99^th^ percentile), white circles the median and gray boxes the interquartile range. Boxplots represent the distribution of the data at the level of individual mice, whiskers the data range, boxes the interquartile range and horizontal line the median. *: p < 0.05, **: p < 0.01, *** p < 0.001. See **Table S1** for all statistical information.

### Adult labeled L5 IT (Tlx3) neurons are anatomically distinct and their activity is less affected by clozapine administration

In attempting to repeat our experiments with AAV-driven GCaMP expression, we found that Cre-dependent labeling in the Tlx3-Cre mouse is strongly developmentally dependent. Indeed, the timing of labeling captured anatomically distinct neuronal subpopulations. The strategy used in the experiments of **Figure 2** was to cross Tlx3-Cre mice with a Cre-dependent reporter line. We refer to these cells as *all* Tlx3 neurons since this labeling strategy permanently tags cells in which Cre recombination events occurred any time between early development and adulthood. An alternative strategy we used was the injection of an AAV expressing a Cre-dependent reporter (EGFP) in adult (> P28) mice to tag neurons that express Cre at the time of injection. We call these cells *adult labeled* Tlx3 neurons. We employed both labeling strategies in the same mice by injecting AAV2/1-Ef1α-DIO-EGFP in primary visual cortex (V1, also known as VISp) of Tlx3-Cre crossed with Ai14 mice that express tdTomato in a Cre-dependent manner (**Figure 3A**, left). We noticed that, among the population of all Tlx3 neurons, there was a set of cells located in upper L5 that were not labeled by AAV driven expression in adult mice (tdTomato positive only, **Figure 3A**). We call these cells *developmentally labeled* Tlx3 neurons. Adult labeled neurons were located deeper in cortex compared to developmentally labeled ones (**Figure 3A**). To test if this asymmetry was an artifact of the AAV tropism or of the delivery method, we repeated the experiment by injecting a different viral serotype (PHP.eB-Ef1α-DIO-EGFP) (**Figure 3B**) and by delivering the AAV through retro-orbital deposits instead of intracranial injections (**Figure 3C**). In all these experiments, adult labeled Tlx3 neurons were located deeper in cortex than developmentally labeled ones (**Figure S2A, B**). These results show that, depending on the developmental timing, the Tlx3-Cre line can tag anatomically different subsets of cells (**Figure 3D**).

**Figure 3.**
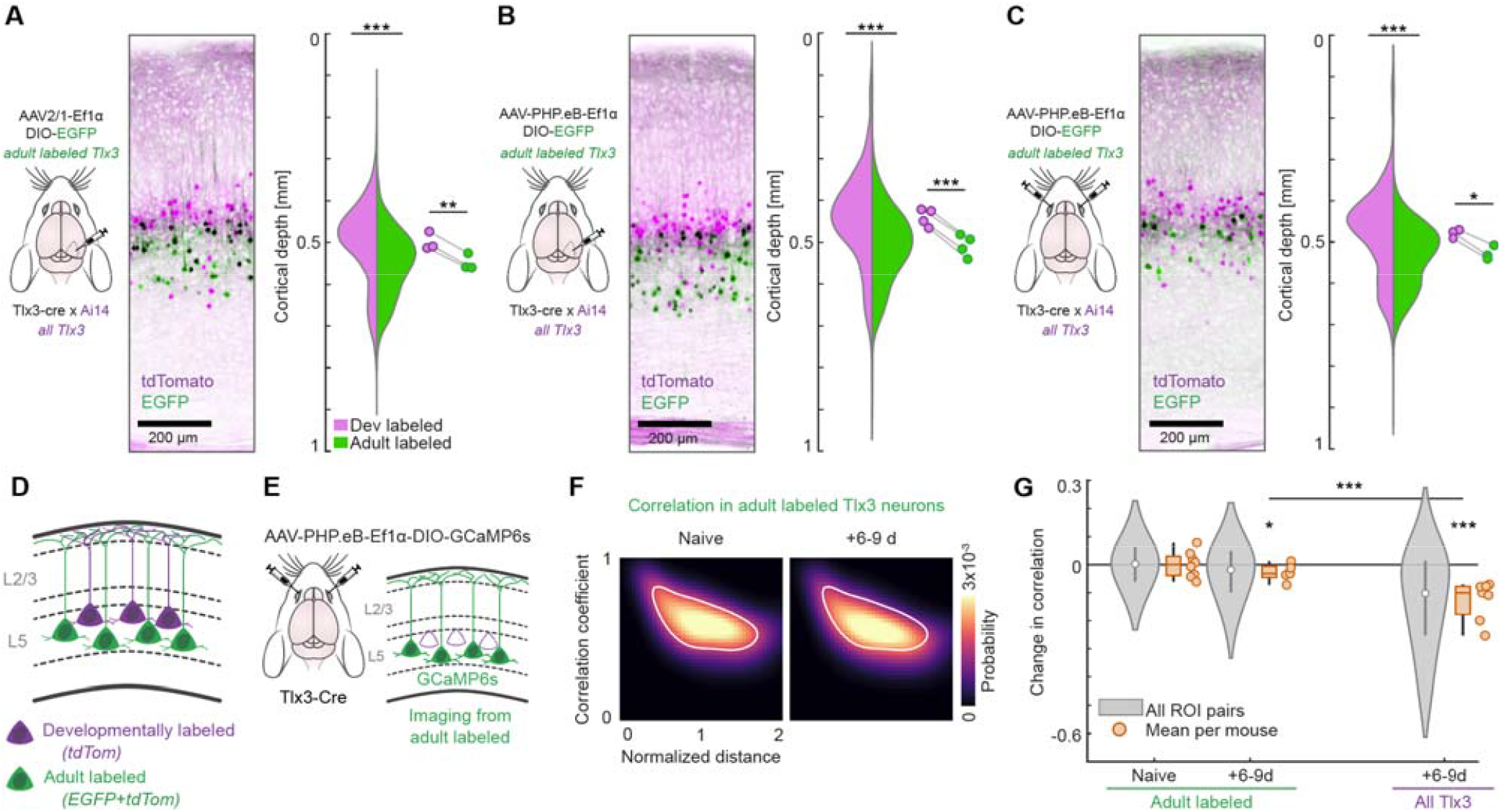
Anatomically distinct L5 IT neurons are differentially affected by clozapine. **(A)**Adult labeled Tlx3 neurons are located deeper in cortex. Left: Schematic of the experiment. Middle: Representative histological image. Right: Depth distribution of labeled neurons in V1, violins represent the distribution of depth for developmentally (magenta) and adult labeled (green) Tlx3 neurons. Data points mark averages by mouse. *: p < 0.05, **: p < 0.01, ***: p < 0.001. See **Table S1** for all statistical information. **(B)**As in **A**, but using a different AAV serotype (PHP.eB). **(C)**As in **B**, but using a different AAV delivery method (bilateral retro-orbital deposit). **(D)**Two, partially overlapping, subgroups of L5 IT Tlx3 positive neurons can be distinguished based on the labeling strategy. Developmentally labeled Tlx3 neurons are tagged by crossing Tlx3-Cre mice with a tdTomato reporter line (Ai14). Adult labeled Tlx3 neurons are tagged by injecting an AAV to drive the expression of a Cre-dependent EGFP reporter. **(E)**Experimental strategy to drive the expression of GCaMP in adult labeled Tlx3 neurons. **(F)**Probability distribution heatmaps of correlation coefficients for all pairs of ROIs as a function of normalized (lambda-bregma) distaance between them. Data recorded from adult labeled Tlx3 neurons. The white contour line represents 50% of the maximum probability of the naive time point. **(G)**Long-term change in correlation for adult labeled Tlx3 neurons (left) with respect to baseline in mice injected with clozapine. As a comparison, data from all Tlx3 neurons (same data as in **Figure 2C**) are included on the right side. Violin plots represent the distribution of correlation changes for all ROIs (from 1st to 99th percentile), white circles the median and gray boxes the interquartile range. Boxplots represent the distribution of the data at the level of individual mice, whiskers the data range, boxes the interquartile range and horizontal line the median.

We tested if adult labeled Tlx3 neurons, a subset of the neurons whose activity we recorded in the experiments of **Figure 2**, would also show decorrelation of activity after administration of a single dose of clozapine. We selectively expressed GCaMP6s in adult labeled Tlx3 neurons through retro-orbital deposit of an AAV (PHP.eB-Ef1α-DIO-GCaMP6s) (**Figure 3E**) and recorded widefield calcium activity across dorsal cortex. We calculated the correlation of activity across ROIs in naive mice, and 6-9 days after clozapine injection. While adult labeled Tlx3 neurons did exhibit a decrease in correlations, this effect was significantly smaller than that observed in all Tlx3 neurons (**Figure 3F, G**). These data indicate that clozapine preferentially decorrelates the activity of developmentally labeled Tlx3 neurons and has smaller effects on adult labeled ones.

### Clozapine-sensitive L5 IT neurons have a distinct functional and molecular profile

Given the differential effects of clozapine on the two subpopulations of Tlx3 neurons, we sought to identify the functional and molecular features that distinguish clozapine-sensitive from clozapine-insensitive L5 IT neurons. Identifying such features could provide insight into the mechanisms through which clozapine alters cortical dynamics and may reveal molecular targets for more selective pharmacological interventions. We performed fluorescence-guided whole-cell patch-clamp recordings from both Tlx3 neuron subsets. Using an approach analogous to that described in **Figure 3**, we labeled all Tlx3 neurons by crossing Tlx3-Cre mice with a tdTomato reporter line and identified adult labeled Tlx3 cells through AAV injection in adult mice. We then prepared acute brain slices containing V1 and targeted L5 neurons for recording based on their fluorescence profiles (**Figure 4A**). Developmentally and adult labeled neurons exhibited comparable resting membrane potential (**Figure 4B**) and membrane capacitance (**Figure S3A**). Interestingly, we observed differences that indicate decreased neuronal excitability in developmentally labeled Tlx3 neurons. These neurons exhibited a significantly smaller input resistance and shorter membrane time constant compared to adult labeled ones (**Figure 4C-F**). We then characterized suprathreshold firing properties (**Figure 4G**). Although action potential amplitude and width did not differ between populations (**Figure S3B, C**), developmentally labeled Tlx3 neurons displayed reduced excitability relative to adult labeled ones. Indeed, injection of depolarizing current steps of increasing amplitude (**Figure 4G**) revealed that developmentally labeled Tlx3 neurons required higher current thresholds to elicit action potentials (**Figure 4H**) and generated fewer action potentials across all stimulation intensities (**Figure 4I**) in comparison to adult labeled ones. We found no evidence of differences in burst firing behavior between the two subpopulations (**Figure S3D-F**). Collectively, these findings indicate that clozapine-sensitive, developmentally labeled Tlx3 neurons exhibit a general pattern of reduced intrinsic excitability compared to adult labeled ones.

**Figure 4.**
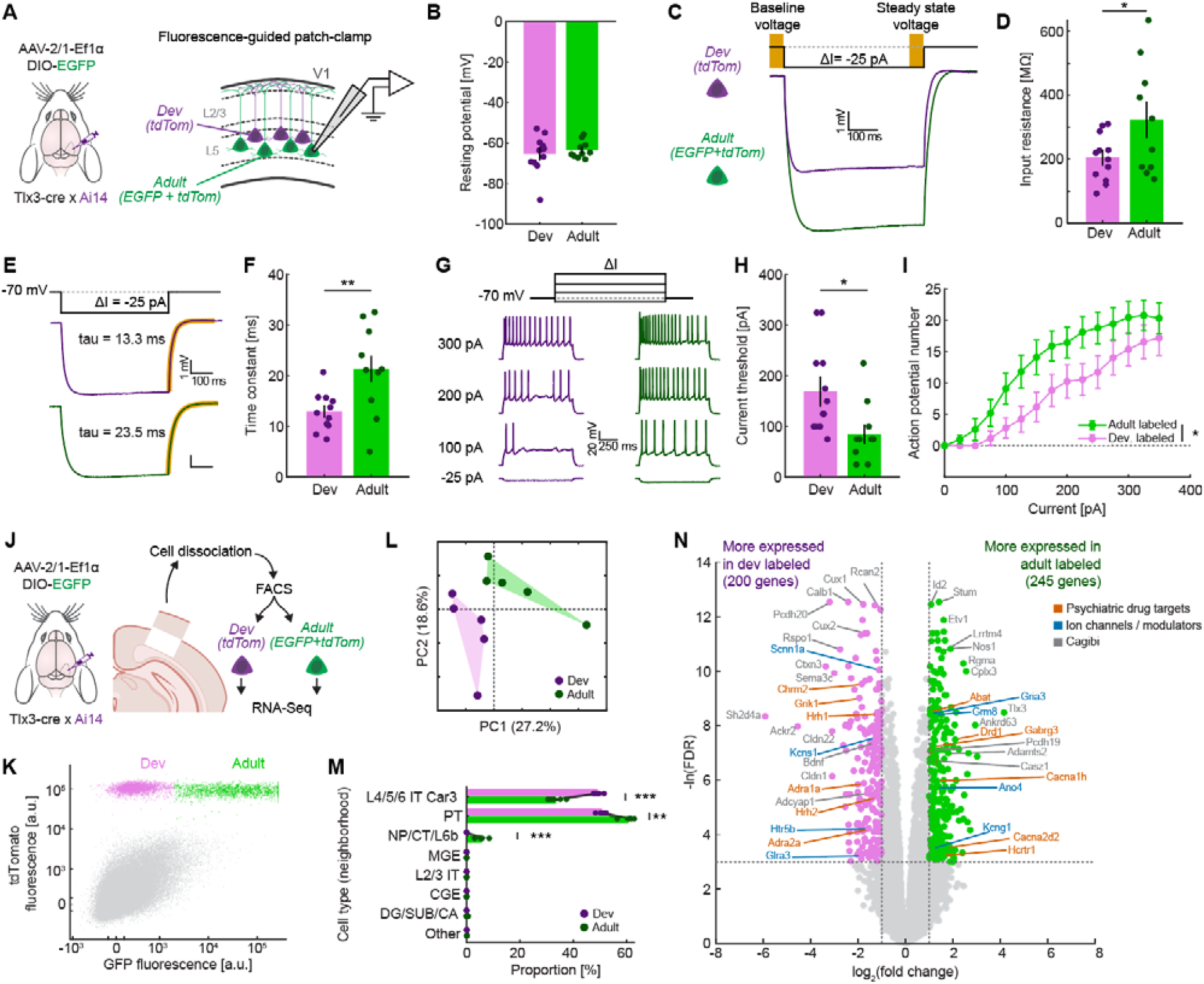
L5 IT neurons that are sensitive to clozapine are a functionally and molecularly distinct population. **(A)**Schematic of the experiment. Left: All Tlx3 neurons were tagged with tdTomato by crossing Tlx3-Cre with Ai14 mice and adult labeled Tlx3 neurons were tagged with EGFP by injecting an AAV in V1. Right: Developmentally labeled Tlx3 neurons are only tagged with tdTomato, while adult labeled neurons are tagged with both tdTomato and EGFP. Acute slices containing V1 were used for fluorescence-guided patch-clamp. **(B)**Resting membrane potential for developmentally labeled and adult labeled Tlx3 neurons. Error bars represent standard error of the mean of the distribution of bootstrapped means, data points represent neurons. **(C)**Example current-clamp traces for developmentally labeled and adult labeled Tlx3 cells showing different input resistance given a small injected current. **(D)**As in **B**, but for input resistance. **(E)**Example current-clamp traces for the measurement of the membrane time constant. A constant current of -25 pA is injected transiently and after the offset of the stimulus the time constant for repolarization (highlighted in orange) is measured. **(F)**As in **B**, but showing the membrane time constant. Here and elsewhere *: p < 0.05, **: p < 0.01, ***: p < 0.001. See **Table S1** for all statistical information. **(G)**Example current-clamp traces of two neurons as a function of injected current. **(H)** As in **B**, but for threshold current to spike. **(I))**Number of action potentials as a function of current, showing excitability for developmentally labeled and adult labeled Tlx3 neurons. Error bars represent standard error of the mean. **(J)**Schematic of the RNA-seq experiment. Left: the labeling strategy is the same as in **A**. Right: the injection site from slices containing V1 was microdissected and individual neurons were dissociated and sorted by fluorescence-activated cell sorting into developmentally labeled neurons (tdTomato-positive) and adult labeled (tdTomato- and EGFP-positive) populations. RNA was then extracted and bulk-sequenced. **(K)** Scatter plot of the cells (for visualization 15% of the data were randomly chosen) used for FACS sorting. Axes are scaled with a biexponential transformation. All detection events were pre-filtered for size, excluding doublets, and excluding high DRAQ7 (dead cells) before being included in the plot. All tdTomato positive cells used for RNA-seq are colored in green and magenta to show adult and developmentally labeled populations. **(L)** PCA on z-scored expression values across all filtered genes. Each point represents a mouse, with convex hulls indicating group boundaries. **(M)**Estimated cell type composition from deconvolution of bulk RNA-seq data for developmentally and adult labeled Tlx3 neurons. Cell type proportions were inferred by deconvolving bulk RNA-seq samples using a single-cell RNA-seq reference atlas from the Allen Institute. Individual data points represent samples, with lines connecting paired samples from the same mouse. IT: intratelencephalic, PT: pyramidal tract, NP: near projecting, CT: cortico-thalamic, MGE: medial ganglionic eminence, CGE: caudal ganglionic eminence, DG: dentate gyrus, SUB: subiculum, CA: cornu ammonis fields. **(N)** Volcano plot of differentially expressed genes between developmentally and adult labeled Tlx3 neurons. Genes that had an absolute fold change larger than 2 and a false discovery rate smaller than 0.05 were colored green (245 genes) if more expressed in adult labeled neurons or magenta (200 genes) if more expressed in developmentally labeled neurons. A curated selection of genes was annotated and categorized as psychiatric drug targets, ion channels and modulators, or general interest. Genes overlapping with the SCHEMA gene set or belonging to SFARI class 1 genes are highlighted with red squares and crosses respectively. This selection is illustrative rather than comprehensive.

We next sought to characterize molecular differences in gene expression between the two populations. Acute slices containing V1 were prepared from mice that had undergone an analogous labeling strategy. Cortical tissue encompassing the injection site was microdissected, and single-cell suspensions were prepared for fluorescence-activated cell sorting (FACS) to separate developmentally and adult labeled Tlx3 neurons based on their distinct fluorescence profiles (**Figure 4J, K**). Total RNA was then purified from the sorted populations and processed for RNA sequencing (RNA-seq) analysis. To assess global transcriptomic differences, we performed principal component analysis (PCA) on expression values across all detected genes (**Figure 4L**). Samples from the two populations were completely separable in the space defined by the first two principal components, indicating distinct overall expression profiles. Hierarchical clustering of sample-to-sample correlations independently confirmed this separation, with all samples correctly grouped by condition (**Figure S4A**). To further validate this finding, we performed cell type deconvolution of bulk RNA-seq samples using a single-cell RNA-seq reference atlas from the Allen Institute (Chu et al. 2022; Yao et al. 2021). This analysis uses bulk expression data to estimate cell type composition in terms of 8 groups of cell subclasses defined in the reference atlas and termed neighborhoods (Yao et al. 2021). As expected, the analysis revealed no significant contributions from glial, inhibitory (MGE, CGE), layer 2/3 (L2/3 IT) and hippocampal (DG/SUB/CA) neighborhoods. Instead, the signal was dominated by infragranular excitatory neighborhoods (L4/5/6 IT Car3, PT, NP/CT/L6b) (**Figure 4M**). Notably, the two populations of Tlx3 neurons exhibited significantly different signatures with very high reproducibility across replicates. Developmentally labeled Tlx3 neurons showed a stronger transcriptomic signature for the L4/5/6 IT neighborhood, whereas adult labeled ones aligned more closely with the PT and NP/CT/L6b neighborhoods. Differential expression analysis identified 445 differentially expressed genes (DEGs; absolute fold change > 2, FDR < 0.05; full list in **Table S3**), with 200 genes enriched in developmentally labeled and 245 genes enriched in adult labeled Tlx3 neurons (**Figure 4N**). Expression of DEGs was reproducible across biological replicates (**Figure S4B, C**). Consistent with differences in laminar distribution, DEGs included upper layer markers (*Cux1, Cux2, Lamp5*) (Nieto et al. 2004; Tasic et al. 2018) enriched in developmentally labeled cells, and the regulator for corticofugal identity *Fezf2* (Arlotta et al. 2005) enriched in adult labeled cells. Furthermore, several DEGs encode targets of approved psychiatric medications, including calcium channels (*Cacna1h*: ethosuximide, *Cacna2d2*: gabapentin, pregabalin), glutamate receptors (*Gria3*: perampanel, *Grik1, Grik3*: topiramate), and modulatory GPCRs (*Drd1*: most AP drugs, *Chrm2*: scopolamine, clozapine, olanzapine, *Hcrtr1*: suvorexant, lemborexant, daridorexant), suggesting potential avenues for biasing pharmacological manipulation towards one of these subpopulations.

Together, these findings demonstrate that developmentally and adult labeled Tlx3 neurons constitute molecularly and functionally distinct subpopulations, providing a basis for dissecting the cell-type-specific mechanisms underlying clozapine’s long-term effects on cortical activity.

### Clozapine increases reliability of long-range intracortical inhibitory functional influence

We found that clozapine alters the correlation structure of activity across dorsal cortex in a subset of L5 IT neurons on a timescale of days. Decorrelation across dorsal cortex could arise from multiple mechanisms. Given that L5 IT neurons are the primary source of long-range cortico-cortical projections, alterations of direct communication between cortical areas might be driving this effect. To test this possibility, we used an experimental setup to measure the functional influence between brain regions before and at different time points after clozapine administration. We used AAVs to express GCaMP in V1 neurons and the excitatory opsin ChrimsonR in a source region projecting to V1. We then optogenetically stimulated superficial axons in V1 while recording calcium responses in both L2/3 and L5 neurons. We refer to this measurement as functional influence.

We first asked if clozapine alters direct cortico-cortical functional influence. We expressed ChrimsonR in cingulate cortex (in either anterior cingulate (ACC) or retrosplenial cortex (RSP)) which provides one of the primary inputs to V1 (Leinweber et al. 2017) and compared functional influence between saline and clozapine injected mice (**Figure 5A**). Consistent with previous results (Leinweber et al. 2017; Zhang et al. 2014), stimulating cingulate axons resulted in net positive responses in V1 (**Figure 5B**). Acutely, at 1 hour after injection, clozapine strongly increased the strength of the functional influence. However, this effect was undetectable 24 hours later. Note, 5 to 9 days after injection, it appeared as if there were a trace of a stimulation-driven decrease in activity in clozapine injected mice, but this was not significant at the population level.

**Figure 5.**
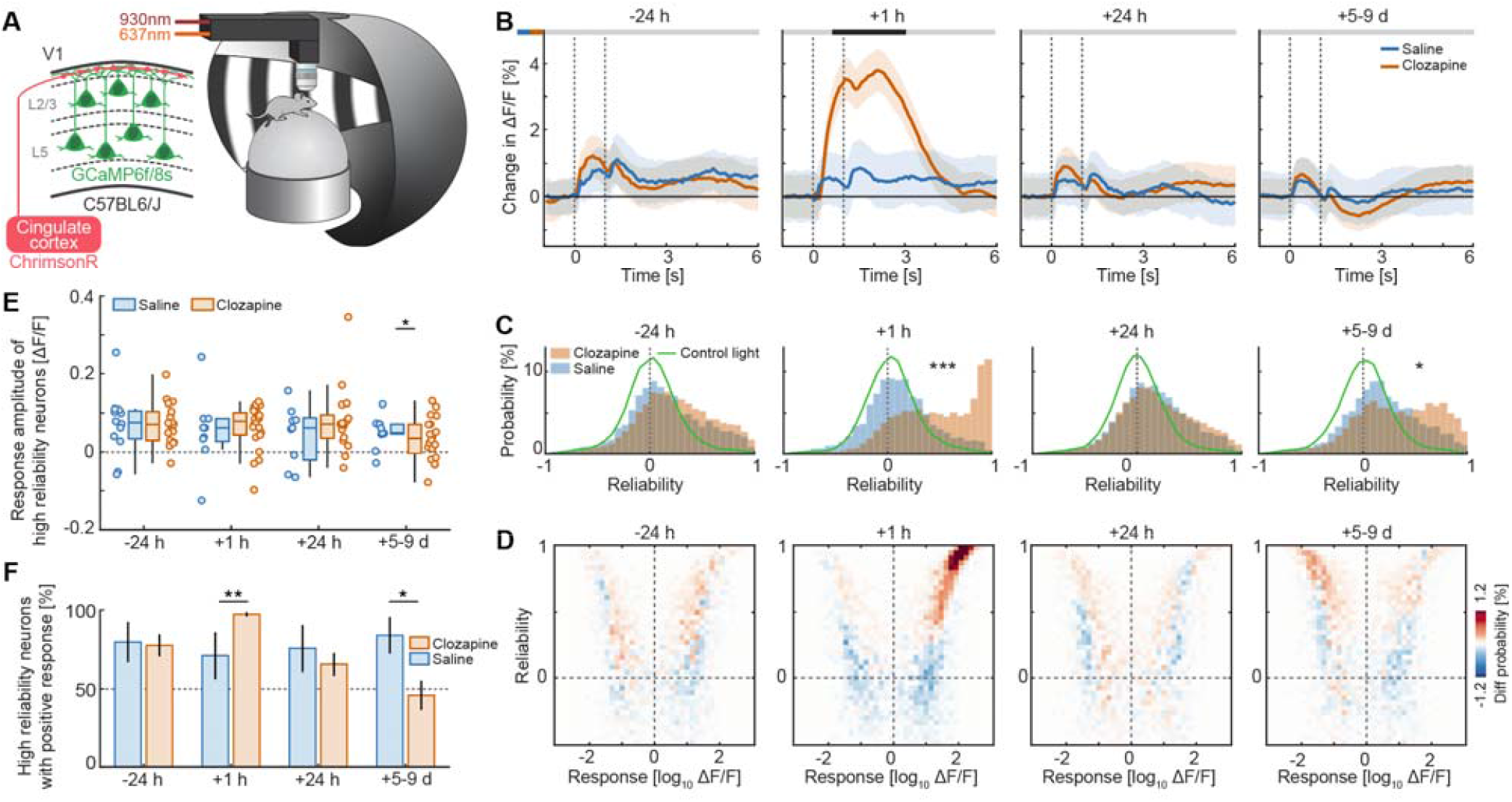
Clozapine increases reliability of long-range intracortical inhibitory functional influence. **(A)**Schematic of the experiment. A GCaMP variant was expressed in V1, and ChrimsonR was expressed in cingulate cortex through AAV injections. In V1, we optically stimulated the long-range axons from cells in cingulate cortex and recorded responses in V1 cells. **(B)**Population responses to the optogenetic stimulation before and at different time points after injection. The shading represents one standard deviation of the hierarchically bootstrapped distribution of means, and the vertical dashed lines the stimulus onset and offset. The horizontal bar above the plot represents time bins in which the responses in saline and clozapine injected mice are statistically different (gray: not significant, black: p < 0.05). See **Table S1** for all statistical information. **(C)**Distribution of reliability of the responses to optogenetic stimulation. The green line represents the kernel density estimate of the responses to a control light stimulation. Here and elsewhere *: p < 0.05, **: p < 0.01, ***: p < 0.001. **(D)**Density plot (40 by 40 bins) showing the difference between the distribution of neurons from saline and clozapine injected mice in the space of reliability and response amplitude. **(E)**Response amplitude of neurons with high reliability (≥ 0.85). Whiskers represent the data range, boxes the interquartile range, and the horizontal line the median. Each point represents data from one mouse. **(F)**Fraction of high reliability neurons with a positive response. Bars and error bars represent the mean and standard deviation of the hierarchically bootstrapped distribution of means.

To investigate this more closely, we measured the reliability of these responses. For each ROI we considered all stimulus repetitions in a session and defined reliability as the correlation coefficient between the average response curves of even and odd trials (**Figure 5C**) (Pachitariu et al. 2018). We used a control light stimulus to estimate the combined influence of non-optogenetic stimulation-driven fluctuations in calcium activity and putative visual stimulation on our reliability measurements. This demonstrated that most neurons did not respond to our optogenetic stimulation beyond what would be expected from the control stimulation distribution (**Figure 5C**, only 18.3% of neurons were above the 97.5^th^ percentile of the control distribution). This is in line with previous reports showing that only about 10% of the neurons in V1 show significant responses to stimulation of axons from ACC (Leinweber et al. 2017). We found that the reliability of responses sharply increased 1 hour after the administration of clozapine, mirroring the increase in amplitude of the responses, but returned to pre-injection levels 24 hours after. Surprisingly, a week after injection the reliability of the responses in clozapine injected mice increased again and was significantly higher than in saline controls.

We then asked if neurons that increase their reliability one week after the injection of clozapine are preferentially functionally excited or inhibited. We created 2D density plots of the distribution of all cells in the space of response amplitude and reliability (**Figure S5A, B**) and plotted the difference between clozapine and saline injected distributions for each time point (**Figure 5D**). Acutely, at 1 hour after injection, the majority of neurons with high reliability had positive responses, consistent with the strong increase in mean response (**Figure 5B**). Strikingly, the opposite was true at 5 to 9 days post injection, where the neurons with high reliability had predominantly negative responses in the clozapine mice. To quantify this, we selected highly reliable neurons (reliability ≥ 0.85) and quantified the average responses of this population. One week after injection, reliable neurons had a significantly lower response in clozapine mice than in saline control mice (**Figure 5E**). Moreover, while in naive mice the majority (∼80%) of reliable neurons responded positively, one week after clozapine injection this proportion flipped and about 60% of reliable neurons had negative responses (**Figure 5F**). These results were robust to different thresholds used to define reliable neurons (**Figure S6A-L**) and selecting cells with low reliability (between -0.5 and 0.5) resulted in no difference between the treatment groups (**Figure S6M-O**).

Taken together, these data indicate that a single dose of clozapine can modulate functional influence across cortical areas and suggest that what we observed as decorrelation of activity across cortical regions could be mediated by the increase in reliability of intracortical inhibitory functional influence.

### Clozapine increases reliability of inhibitory thalamo-cortical functional influence from higher-order but not first-order thalamus

An additional circuit-level alteration that could account for the decorrelation of the L5 IT network, beyond changes to direct interactions, is a change to cortico-thalamic and cortico-striatal-thalamic loops. To begin characterizing this, we measured the clozapine-induced changes to thalamo-cortical functional influence. We used AAVs to express ChrimsonR in thalamus and GCaMP in V1 (**Figure 6A**). The thalamic injections were aimed at first-order and higher-order visual nuclei of thalamus. Note, expression was typically not confined to just one thalamic nucleus. For analysis, we split mice into two groups, those with predominantly first-order thalamus labeling, and those with predominantly higher-order thalamus labeling, based on histological analysis (**Figure 6B**). For simplicity we will refer to these groups simply as labeled in first-order or higher-order thalamus. The optogenetic stimulation in the two groups of mice resulted in very different responses in V1: Stimulating the axons from first-order thalamus elicited an increase in calcium activity, while stimulating the ones from higher-order thalamus did not, as previously reported (Fang et al. 2020; Furutachi et al. 2024) (**Figure 6C, E**). A single threshold applied on early response amplitudes prior to treatment separated first-order from higher-order thalamus-labeled mice (**Figure S7**).

**Figure 6.**
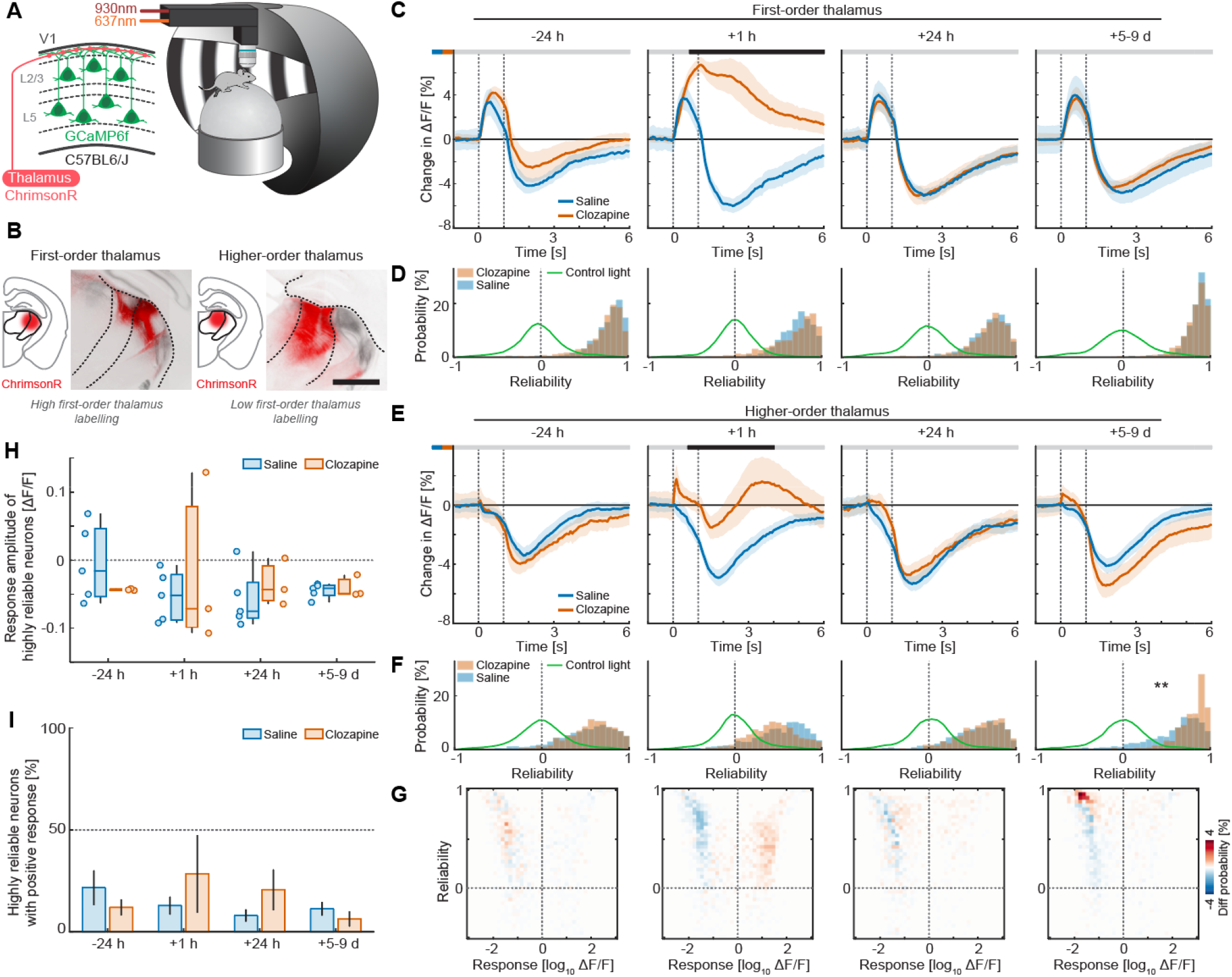
Clozapine increases the reliability of higher-order thalamo-cortical inhibitory influence. **(A)**Schematic of the experiment. A GCaMP variant was expressed in V1, and ChrimsonR was expressed in thalamus through AAV injections. In V1, we optically stimulated the long-range axons from cells in thalamus and recorded responses in V1 neurons. **(B)**Experimental mice were classified into two groups: first-order thalamus and higher-order thalamus based on the amount of ChrimsonR expression in the two areas. Scale bar: 1 mm. **(C)**Population responses to the optogenetic stimulation of first-order thalamus axons before and at different time points after injection. The shading represents one standard deviation of the hierarchically bootstrapped distribution of means, the vertical dashed lines the stimulus onset and offset. The horizontal bar above the plot represents time bins in which the responses in saline and clozapine injected mice are statistically different (gray: not significant, black: p < 0.05). See **Table S1** for all statistical information. **(D)**Distribution of reliability of the responses to optogenetic stimulation of first-order thalamus axons. The green line represents the kernel density estimate of the responses to a control light stimulation. **(E)**As in **C**, but for stimulation of higher-order thalamus axons. **(F)** As in **D**, but for stimulation of higher-order thalamus axons. *: p < 0.05, **: p < 0.01, ***: p < 0.001. **(G)**Density plot (40 by 40 bins) showing the difference between the distribution of neurons from saline and clozapine injected mice in the space of reliability and response amplitude. **(H)**Response amplitude to the stimulation of higher-order thalamus axons of neurons with high reliability (≥ 0.85). Whiskers represent the data range, boxes the interquartile range, and the horizontal line the median. Each data point is one mouse. **(I)**Fraction of high reliability neurons with a positive response to the stimulation of higher-order thalamus axons. Bars and error bars represent the mean and standard deviation of the hierarchically bootstrapped distribution of means.

Acutely, at 1 hour after clozapine injection, we found a strong increase in the functional influence of first-order thalamus on V1. By 24 hours, and at 5 to 9 days after injection, the functional influence was not different from saline controls (**Figure 6C**) with no detectable differences in reliability at any time point (**Figure 6D**). For higher-order thalamus, one hour after injection, the inhibitory component was largely abolished, mirroring the acute increase of functional influence that we observed for cortico-cortical connections. This effect resolved by 24 hours, and at 5 to 9 days post injection we only observed a modest, not statistically significant, decrease of mean response amplitude (**Figure 6E**). The reliability of responses evoked by stimulation of axons from higher-order thalamus remained unchanged up to 24 hours after clozapine injection. Interestingly, however, we saw an increase in reliability 5 to 9 days after clozapine administration (**Figure 6F**). To determine whether the neurons with increased reliability display positive or negative responses, we plotted the difference of the 2D distributions of reliability and response amplitude between clozapine and saline injected mice. While clozapine acutely shifted responses towards positive values, the 5 to 9 day time point showed an increase in reliability specifically among negatively responding neurons (**Figure 6G**), suggesting that a week after clozapine administration higher-order thalamus maintains an inhibitory functional influence on cortex but it does so with increased reliability. If this interpretation is correct, one would expect the response sign and amplitude of highly reliable cells to not change as a function of the treatment. Consistent with this prediction, we found no evidence of differences in either mean response amplitude (**Figure 6H**) or the fraction of positively responding neurons (Figure 6I) among highly reliable cells between saline and clozapine injected mice at any time point.

Thus, our data indicate that clozapine preferentially affects the influence of the higher-order thalamus on cortex at 5 to 9 days post injection. These effects manifest as an increase in the reliability of functional inhibition, mirroring the patterns seen in long-range cortico-cortical influence.

### Clozapine does not have long-term effects on visual responses

Clozapine has been reported to alter sensory event-related potentials in patients with schizophrenia, with effects that emerge over weeks to months of treatment (Nagamoto et al. 1999; Pallanti et al. 1999; Umbricht et al. 1998). These findings suggest that clozapine can induce long-term changes in how sensory information is processed in the brain. Given these observations, we asked whether a single dose of clozapine is sufficient to alter basic features of visual sensory processing in mice. To test this, we compared V1 responses to full field sinusoidal drifting gratings following a single clozapine injection. We used sequences of brief (3 s) presentations of gratings drifting in a randomly chosen direction (of 8 possible directions; **Figure 7A**). In saline injected mice, individual neurons exhibited a response profile that was similar across sessions (**Figure 7B, D**). In clozapine mice, we again found a strong transient increase in responses at 1 hour after injection, which returned to baseline by 24 hours and did not change thereafter (**Figure 7C, D**). Consistent with this, we only found acute effects on response reliability: Clozapine acutely increased reliability at 1 hour after injection. This effect, however, dissipated by 24 hours, with no detectable differences between clozapine and saline injected mice at any subsequent time point (**Figure 7E, S8**).

**Figure 7.**
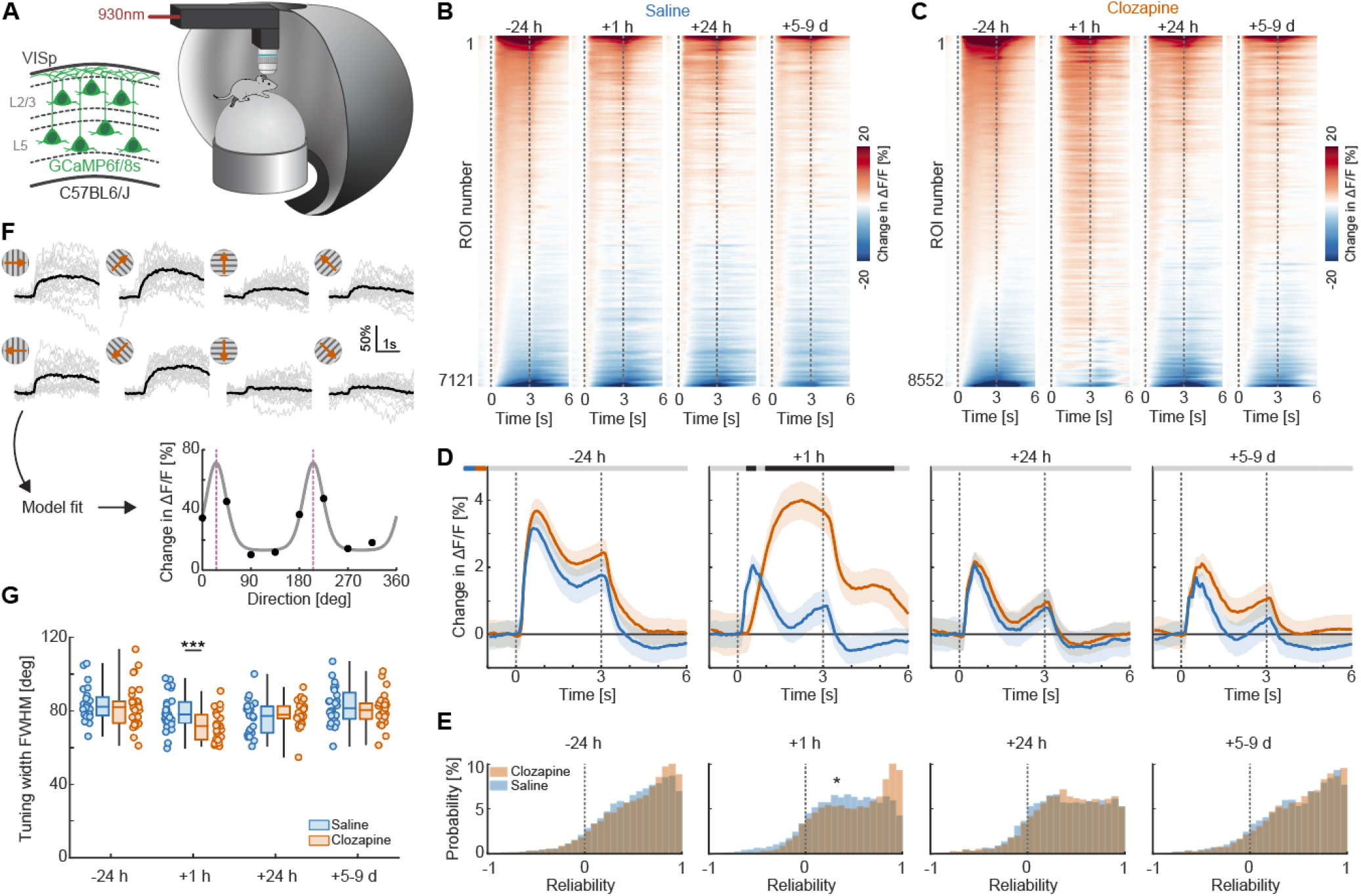
Clozapine does not have long-term effects on bottom-up visual inputs. **(A)**Schematic of the experimental setup. We recorded responses to full field drifting sine gratings in V1. **(B)**Heatmap of responses to grating onset of all chronically recorded neurons in saline injected mice. Neurons were sorted based on the response amplitude of the first time point (-24 h) and the sorting order was kept for all the following time points. To avoid graphical aliasing the heatmaps are smoothed vertically (only across neurons) with a Gaussian kernel. **(C)**As in **B**, but for all chronically recorded neurons in clozapine injected mice. **(D)**Average population response to gratings of all recorded neurons. The shading represents plus and minus one standard deviation of the hierarchically bootstrapped distribution of means, the vertical dashed lines the stimulus onset and offset. The horizontal bar above the plot represents time bins in which the responses in saline and clozapine injected mice are statistically different (gray: not significant, black p < 0.05). See **Table S1** for all statistical information. **(E)**Distribution of reliability of the responses to gratings. Here and elsewhere *: p < 0.05, **: p < 0.01, ***: p < 0.001. **(F)**Top: responses of an example orientation selective neuron to 8 grating directions. Gray curves represent responses from all time points, and the black curve represents the mean response. Bottom: tuning curve fit for that neuron, black dots represent the measured responses, the pink dotted line the preferred orientation of the fit. **(G)**Mean tuning width (full-width half-maximum) of the orientation selective neurons for each mouse. Only mice with at least 5 orientation selective neurons were included in the analysis. Whiskers represent the data range, boxes the interquartile range, and the horizontal line the median. Each data point is one mouse.

To test whether we could find evidence of any more subtle changes to the visual responses, we characterized orientation selectivity. We fit a tuning curve model (see **Methods**) to each neuron’s responses across the 8 grating directions (**Figure 7F**). For each neuron, we then determined whether the response was orientation selective (**Methods**) and restricted further analysis to selective neurons. The average tuning width of orientation-selective neurons decreased 1 hour after clozapine treatment relative to saline controls; however, no differences between treatment groups were detected at any subsequent time point (**Figure 7G**).

In sum, we find no evidence of clozapine-induced long-term effects on visual responses in V1. This is consistent with the observation that functional influence from first-order thalamus, one of the main drivers of feedforward sensory signals in V1, was not significantly altered by clozapine treatment. These results suggest that the long-term circuit alterations driven by clozapine are specific to long-range intracortical circuits and loops involving higher-order thalamus, rather than affecting primary sensory pathways.

### Long-term decorrelation is shared by AP drugs with different receptor profiles

If the long-term effects we observed are relevant to AP action, rather than a peculiarity of clozapine, they should be shared by AP drugs with different pharmacological profiles. Thus, a key question is whether the long-term functional effects of a single dose are specific to clozapine or a common feature of AP drugs. We focused on the decorrelation of L5 IT network activity because it is the network-level phenotype that provides an integrated readout of the long-term circuit changes we describe. To address this question, we repeated the experiments of Figure 2 with three other AP drugs: aripiprazole, a partial agonist at D2 receptors; risperidone, an antagonist at both serotonin 5-HT2A and D2 receptors; and haloperidol, a potent D2 receptor antagonist with little affinity for serotonin receptors (Figure 8A). A single dose of each of the three AP drugs resulted in a decorrelation of L5 IT network activity 4 to 9 days after injection that was significantly larger than in saline injected mice (Figure 8B-C). Thus, despite their different pharmacology, all AP drugs we tested share a long-term effect on L5 IT network activity that is visible many days after a single treatment, suggesting that the circuit changes we describe for clozapine may be a common feature of AP drug action.

**Figure 8.**
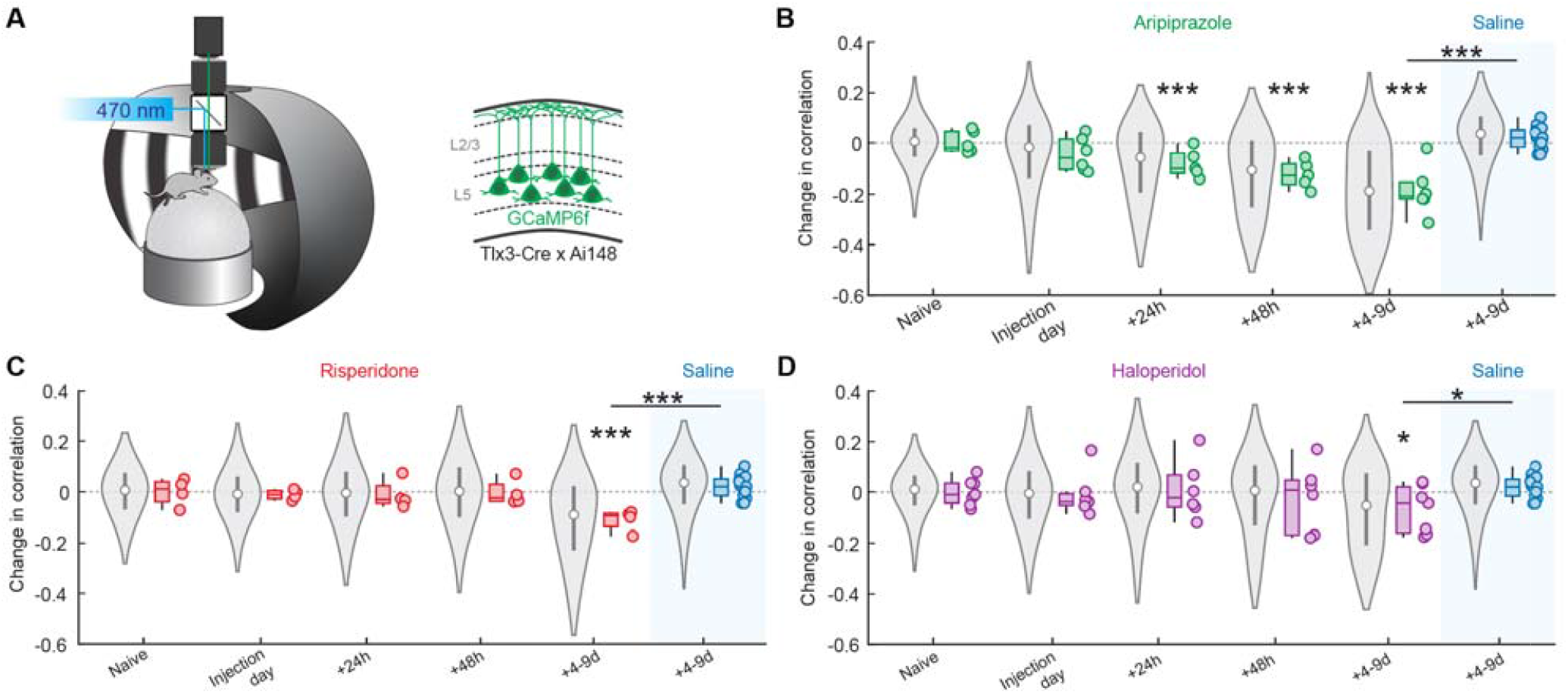
Antipsychotics with different pharmacology induce long-term decorrelation in the activity of a subset of L5 IT neurons. **(A)**Left: Schematic of the experimental setup. Right: GCaMP6f was expressed in a subset of L5 IT neurons by crossing Tlx3-Cre with Ai148 mice. **(B)**Time course of the change in correlation with respect to baseline in aripiprazole injected mice. Long-term change in correlation in saline treated mice is shown for comparison (same data in B-D). Violin plots represent the distribution of correlation changes for all ROIs (from 1st to 99th percentile), white circles the median and gray boxes the interquartile range. Boxplots represent the distribution of the data at the level of individual mice, whiskers the data range, boxes the interquartile range and horizontal line the median. *: p < 0.05, ***: p < 0.001. See **Table S1** for all statistical information. **(C)**As in B but for mice injected with risperidone. **(D)** As in B but for mice injected with haloperidol.

## DISCUSSION

In this study we show that a single dose of clozapine, the only approved medication for treatment-resistant schizophrenia (Agid et al. 2024; Flanagan et al. 2020), is sufficient to modulate behavior, as well as cortical dynamics and long-range communication for up to 9 days after administration in mice. Specifically, we found long-term alterations in several aspects of spontaneous behavior that paralleled the time course of the progressive decorrelation of activity in a genetically defined subset of L5 IT neurons. We functionally and molecularly characterized this clozapine-sensitive subpopulation and provide evidence of two possible circuit alterations that could underlie the decorrelation. We believe that these results constitute both a promising starting point to begin deciphering the circuit mechanism of action of clozapine and a proof of principle for previously underappreciated long-term effects of the drug.

The therapeutic efficacy of clozapine typically builds over days to weeks (Agid et al. 2003). Although factors such as slow dose titration in patients and limited sensitivity of clinical assessments may contribute to this delay, this timescale far exceeds the pharmacokinetic lifetime of the drug, which is estimated to be on the order of hours in both plasma and brain (Baldessarini et al. 1993; Jendryka et al. 2019). This temporal dissociation, together with our series of observations of long-term effects following acute administration, suggests that clozapine drives targeted plasticity sufficient to induce sustained alterations in neural circuit function, rather than relying solely on acute receptor occupancy. Based on this, we propose that less-than-daily dosing regimens warrant more thorough clinical exploration.

### Long-term effects and neuroplasticity

A temporal dissociation between drug action and realization of full improvements is not unique to AP drugs. A similar logic applies to antidepressants, where early reports (Vetencourt et al. 2008) and more recent findings (Castrén and Antila 2017) suggest that the drugs acutely act as a trigger for slower synaptic and structural plasticity. Similarly, clozapine might work by initiating a cascade of events, likely involving neuronal plasticity, that gradually reshapes circuit dynamics (Konradi and Heckers 2001). If this is the case, two fundamental questions arise.

The first concerns specificity: Changes required to ameliorate psychotic symptoms are presumably small and circuit-specific, yet clozapine has a broad pharmacology (Coward 1992; Roth et al. 2004) and distributes throughout the brain. How can a low-dimensional intervention produce what are likely very specific circuit effects? One possibility is that the drug’s specificity comes from its promiscuity: The simultaneous engagement of multiple receptor systems across different cell types could produce combinatorial modulation with specific functional consequences (Roth et al. 2004). Alternatively, although the pharmacological signal is diffuse, the capacity for plasticity might be constrained by cell-type- or region-specific structural and molecular features. Resolving the relative contribution of these non-mutually exclusive mechanisms will require a systematic and comparative assessment of plasticity in different cell types and brain regions in response to pharmacologically diverse AP drugs.

The second question concerns the identity of the circuits that undergo plasticity. This study offers a preliminary entry point: we found that clozapine produced a lasting increase in the reliability of inhibitory functional influence in both cortico-cortical and thalamo-cortical pathways. Since the vast majority of both intratelencephalic and thalamo-cortical projections are glutamatergic, this inhibitory influence must be mediated by local inhibitory neurons in cortex. Whether the relevant site of plasticity is the synapse from excitatory projection neurons onto local inhibitory cells, the synapse from inhibitory cells onto neighboring excitatory neurons, or both, remains to be determined.

### Cortical signatures of antipsychotic efficacy

While earlier accounts of psychosis have focused on subcortical dopamine dysregulation (Howes and Kapur 2009; Schmack et al. 2021) or cortical glutamatergic synaptopathy (Heinz et al. 2019; Howes and Shatalina 2022; Howes and Onwordi 2023; Lisman et al. 2008), there is growing recognition that dysfunction at the level of cortical circuit dynamics may be a common feature across these frameworks. Specifically, computational approaches to psychosis posit that alterations to cortical activity patterns driven by changes in long-range communication may play a central role in the pathophysiology (Keller and Sterzer 2024; Sterzer et al. 2018). For example, heightened strength of lateral communication in cortex could be a key driver of psychosis. Consistent with this, multiple AP drugs have been shown to produce a decorrelation of activity in deep cortical layers (Heindorf and Keller 2024). Our results now further expand these findings: the decorrelation persists for over a week after a single dose and is accompanied by changes in cortico-cortical and thalamo-cortical functional connectivity consistent with a reduction in long-range L5 communication. These findings identify, for the first time, specific long-range circuits that are durably modulated by AP drugs. They corroborate the idea that cortical communication is central to both psychosis and its treatment, and open the intriguing possibility that treatment-effective plasticity in cortical circuits could be induced through alternative pharmacological strategies that do not rely on the classic D2/5-HT2A axis.

### Dosing and adverse effects

Clozapine is the only medication recommended for treatment-resistant schizophrenia, but its use is limited by severe adverse effects (Flanagan et al. 2020), several of which are related to plasma concentration (Skokou et al. 2022). Dosing strategies to reduce exposure to the drug while maintaining therapeutic efficacy could therefore lower the risk of adverse effects. Direct evidence for other AP drugs is limited to a single trial and suggests this is a viable approach: In stabilized patients, taking AP drugs every second day was not associated with a higher risk of symptom exacerbation or relapse than daily dosing (Remington et al. 2011). Similarly, long-acting injectable risperidone can maintain clinical stability without sustained D2 receptor occupancy at the levels associated with acute efficacy (Uchida et al. 2008; Ikai et al. 2012; Uchida and Suzuki 2014). For clozapine, such strategies have not been tested, and there are specific reasons for caution. Abrupt interruption of clozapine treatment can cause withdrawal symptoms and, after more than 48 h without the drug, re-titration is cautiously recommended to not lose tolerance to the hypotensive effect of the drug (U.S. Food and Drug Administration 2025). However, these risks have been documented for the interruption of an established daily regimen, and it is unclear to what extent they would apply to a regimen designed with longer dosing intervals from the start of treatment. Moreover, the 48-hour threshold is largely arbitrary and the time over which tolerance is lost is not known (Flanagan and Every-Palmer 2025). Our results are not to be translated directly into a dosing regimen. However, based on our results, we speculate that if the therapeutically relevant effects of AP drugs develop slowly and depend on plasticity, the frequency of dosing may matter less than currently assumed, and alternative dosing strategies may warrant further clinical exploration.

### Limitations of the study

Our study has three major caveats:

1. We do not know whether any of the effects caused by clozapine we describe are treatment-relevant. We suspect, however, that they are, based on three arguments. First, the time course of effects roughly matches the comparably slow onset of therapeutic effectiveness. Second, the effects involve long-range communication in cortex that is also thought to be essential in the etiology of schizophrenia. Third, an increase in long-range functional inhibition is one possibility to reduce spurious activation of cortical networks that drive hallucinatory percepts.
2. Expression of a calcium-binding protein (GCaMP) in a subset of neurons throughout development, as we do for developmentally labeled Tlx3 neurons, could sensitize these neurons to the effects of clozapine (**Figures 2** and **3**). Based on two considerations, however, we speculate that any such effect is unlikely to fully explain our findings. First, physiological and gene expression differences that could explain a differential response to clozapine are also present in the absence of developmental GCaMP expression (**Figures 3** and **4**). Second, the changes in cortico-cortical and thalamo-cortical functional influence that we describe here are measured without developmental expression of GCaMP and are consistent with a decorrelation effect. Testing this possibility will require targeting GCaMP expression to developmentally, but not adult, labeled Tlx3 neurons in adulthood.
3. Our functional influence experiments are not constrained to measuring the influence of developmentally labeled Tlx3 neurons. Specifically, in the case of the cortico-cortical experiments (**Figure 5**), ChrimsonR is expressed unselectively in cingulate cortex. While most long-range input to V1 likely originates from L5 IT neurons in cingulate, we do not know the fraction of developmentally labeled Tlx3 neurons in this population. More broadly, even if we could resolve this question, it would not fully explain our findings, as we lack a complete understanding of the mechanisms that link a general increase in the reliability of long-range inhibition to a preferential decorrelation of developmentally labeled L5 IT neurons. It is conceivable that if we could measure the long-range influence just within the network of developmentally labeled Tlx3 neurons, we would find larger effects. Alternatively, it is possible that the network of developmentally labeled Tlx3 neurons is more sensitive to a general increase in long-range inhibition, as a larger fraction of their synapses are long-range connections.

## ACKNOWLEDGMENTS

We thank all the members of the Keller lab for discussion and support, and Tingjia Lu (FMI vector core) for producing the viral vectors. We also thank the FMI facilities for experimental support: Laure Plantard and Jan Eglinger for anatomical image acquisition, Hubertus Kohler for flow cytometry, the FMI functional genomics facility for RNA sequencing, Hans-Rudolf Hotz for RNA-seq analysis, the FMI media kitchen for assistance with solution preparation, and Jennifer Gröli for mouse husbandry. This project has received funding from the Swiss National Science Foundation (GBK), the Novartis Research Foundation, EMBO postdoctoral fellowship ALTF 844-2022 (LL), and the European Research Council (ERC) under the European Union’s Horizon 2020 research and innovation programme (grant agreement No 865617) (GBK).

## DATA AVAILABILITY

Software for controlling the two-photon and widefield microscopes and preprocessing of calcium imaging data is available at https://sourceforge.net/projects/iris-scanning/. Raw data and code to generate all figures will be deposited in public repositories upon publication.

## AUTHOR CONTRIBUTIONS

LL designed the study, performed behavioral, two-photon imaging, histology, and RNA-seq experiments, and analyzed the data from these, as well as those from widefield imaging experiments; MH and TS performed the widefield imaging experiments; SK performed and analyzed patch-clamp experiments; AS performed behavioral experiments. All authors wrote the manuscript.

## DECLARATION OF INTERESTS

The authors declare no competing financial interests.

## METHODS AND MATERIALS

### Mice

All animal procedures were approved by and carried out in accordance with guidelines of the Veterinary Department of the Canton Basel-Stadt, Switzerland. We used the following mouse lines: C57BL/6J, Tlx3-Cre (Gerfen et al. 2013) to drive the expression of Cre in L5 IT cortical neurons, Ai14 (Madisen et al. 2010) to drive the Cre-dependent expression of tdTomato, and Ai148 (Daigle et al. 2018) to drive the Cre-dependent expression of GCaMP6f. See **Table S2** for the number of mice used for each experiment. The experiments were conducted on a mixture of male and female mice. In female mice, we did not time experiments to a specific phase of the estrous cycle. Mice were group-housed, up to 5 mice per cage, in conventional EU Type 2L cages containing bedding, nesting material and a handling tunnel, and had access to food (Altromin 1328) and water ad libitum. Clozapine (1 mg/kg or 10 mg/kg, diluted in saline and acetic acid), aripiprazole (0.2 mg/kg), risperidone (0.5 or 2.5 mg/kg) or haloperidol (0.1 mg/kg) were injected intraperitoneally. Treatment allocation was randomized within each cage, and injections were stratified by cage.

### Open Field Test

The test was performed in a circular arena (44 cm diameter, 40 cm tall) with opaque walls bearing no visual cues. The floor and the walls of the arena were cleaned with 10% ethanol before each trial. The illumination consisted of a dim LED (M660L3 Thorlabs, 660 nm) and two infrared LEDs to enable video recordings. Video data for the entire session were collected with an overhead grayscale camera (BFS-U3-16S2M) at a temporal resolution of 30 Hz and a spatial resolution of 1080 by 1080 pixels (0.52 mm/pixel). The data were acquired using custom LabVIEW software. Each mouse was habituated to the experimenter by daily handling for 3 to 5 days before starting the test. The same experimenter performed all habituation and test sessions. The length of each session was 11 minutes (including the time for placing and removing the mouse from the arena).

### Behavioral data analysis

The X and Y positions of individual body parts (**Figure 1A**) were extracted from behavioral videos using DeepLabCut (Mathis et al. 2018). Data were trimmed in time to exclude the initial and final segments of the videos when the mouse was placed and recovered from the arena. Total distance traveled and time spent in the center were calculated using the nose coordinates.

To cluster behavioral syllables we fit a Keypoint-MoSeq (Weinreb et al. 2024) model using all body parts except tail-midpoint and tail-tip. Due to memory constraints, we fit the model using data from a subset of videos consisting of 70 (of 150 total) randomly selected videos, excluding those recorded 1 h after clozapine (10 mg/kg) administration when mice were mostly sedated and stationary for the entire duration of the test. This trained model was then applied to cluster behavioral syllables on the entire dataset. Syllables with an overall frequency smaller than 0.5% were excluded from further analysis, resulting in a total of 24 behavioral syllables. For each session, we calculated a transition matrix as the number of transitions between syllables normalized by the total number of transitions. To statistically compare transition matrices between conditions we computed the L1 distance between the average matrix for each condition and compared it with a null distribution of distances between group averages obtained by randomly shuffling the condition labels 10 000 times. To represent behavioral structure in a lower-dimensional space, syllable frequency vectors and flattened transition matrices were embedded in two dimensions using t-SNE (**Figure 1**).

### Surgery and viral injections

For all surgical procedures, mice were anesthetized using a mixture of fentanyl (0.05 mg/kg; Actavis), midazolam (5.0 mg/kg; Dormicum, Roche), and medetomidine (0.5 mg/kg; Domitor, Orion). After surgery, anesthesia was antagonized by a mixture of flumazenil (0.5 mg/kg; Anexate, Roche) and atipamezole (2.5 mg/kg; Antisedan, Orion Pharma). Both were injected intraperitoneally. Analgesics were applied perioperatively on the scalp (2% lidocaine gel), subcutaneously under the scalp (lidocaine 10 mg/kg and ropivacaine 3 mg/kg) and postoperatively (Ethiqa XR, 3.25 mg/kg). At the start of the surgery, eyes were covered with ophthalmic gel (Virbac Schweiz AG).

For widefield imaging experiments, we implanted crystal skull cranial windows (Kim et al. 2016), as previously described (Heindorf and Keller 2024). Briefly, the perimeter of the skull plate overlying cortex was thinned using a dental drill until it could be removed with forceps. Prior to removing the skull plate overlying dorsal cortex, we recorded the location of bregma relative to other landmarks on the skull. Superglue (Pattex) was used to glue the crystal skull in place. The remaining exposed surface of the skull was covered with cyanoacrylate adhesive (Histoacryl, B. Braun) and a custom titanium head bar was fixed to the skull using dental cement (Paladur, Heraeus Kulzer). In a subset of mice, we also deposited a viral vector based on the PHP.eB capsid (Chan et al. 2017) retro-orbitally to drive the expression of GCaMP.

For two-photon imaging experiments, we implanted 4 mm cranial windows. First, we performed a craniotomy above V1 centered 2.5 mm lateral and 0.5 mm anterior to lambda. The dura mater was removed, and the tissue was always kept moist using 0.22 μm-filtered artificial cerebrospinal fluid (ACSF) (126 mM NaCl, 2.5 mM KCl, 1.2 mM NaH_2_PO_4_, 15 mM NaHCO_3_, 10 mM HEPES, 1.2 mM MgCl_2_, 2.4 mM CaCl_2_, 10 mM glucose). Using a glass capillary, we injected viral vectors to drive the expression of either a calcium indicator or a fluorescent reporter in V1 (3-4 injections 100-250 nL each). A stacked glass coverslip assembly (4 and 5 mm coverslips glued with optical adhesive) was then implanted on the skull using dental cement (Croci et al. 2023). We found this approach facilitated precise positioning within the craniotomy, as the physical contact between the larger coverslip and skull surface provided a stable mechanical reference during implantation. A custom titanium head bar was fixed to the skull using Superglue (Pattex) and dental cement (Paladur, Heraeus Kulzer). Care was taken to cover all the exposed surface of the skull in this process.

For intracortical injections of viral vectors outside of a craniotomy, we drilled small craniotomies over the injection location. Injections (100-250 nL each) were performed using a glass capillary. The craniotomies were covered with Superglue (Pattex) and, in cases in which no windows were implanted, we closed the scalp with surgical sutures. To target RSP, we performed two injections, one at 400 μm lateral, 500 μm anterior from lambda, and one at 400 μm lateral, 1.1 mm anterior from lambda, both at a depth of 350 μm from the dura. To target ACC, we performed one injection at 400 μm lateral, 500 μm anterior from bregma at a depth of 850 μm from the dura. To target thalamus, we performed one injection at 1.6 mm lateral, 2.1 mm posterior from bregma at a depth of 2.45 mm from the dura.

### Virtual reality setup and stimulus design

During two-photon and widefield imaging experiments, mice were head-fixed in a virtual reality setup (Leinweber et al. 2014) and free to locomote on a spherical, air-supported treadmill. A visual scene was projected (Samsung SP-F10M) onto a toroidal screen positioned in front of the mouse covering a field of view of approximately 160 degrees horizontally and 100 degrees vertically. The virtual reality setup was configured for one of the following paradigms: Closed loop, in which the visual stimulus consisted of a virtual corridor containing sinusoidal vertical gratings on the walls and a uniform gray floor. In this paradigm, movement in the virtual corridor was coupled to the locomotion speed of the mouse. Open loop, in which a replay of a previous closed loop session, uncoupled to locomotion, was projected. Dark, in which the projector was turned off, although residual dim ambient lighting might have been present. In addition to these paradigms, we also presented full field drifting sinusoidal gratings (8 directions in randomized order, approximately 0.07 cycles/deg spatial frequency, 2 Hz temporal frequency). Each stimulation lasted 3 s with a randomized inter-trial interval between 3 and 7 s during which a uniform gray screen was shown.

### Two-photon imaging

Functional two-photon calcium imaging was performed using a modified commercial microscope (Bergamo, Thorlabs) (Leinweber et al. 2014). The illumination source was a tunable femtosecond laser (Insight, Spectra-Physics), tuned to 930 nm. Emission light was band-pass filtered using a 525/50 nm filter for GCaMP and a 607/70 nm filter for tdTomato (Semrock) and detected using two GaAsP photomultipliers (H7422PA-40, Hamamatsu). Photomultiplier signals were amplified (DHPCA-100, Femto), digitized (NI5772, National Instruments) at 800 MHz, and band-pass filtered around 80 MHz using a digital Fourier-transform filter implemented in custom-written software on an FPGA (PXIe-7965, National Instruments). The scanning system of the microscope was based on a 12 kHz resonant scanner (Cambridge Technology). Images were acquired at a resolution of 750 by 400 pixels (60 Hz frame rate), and a piezo-electric linear actuator (P-726, Physik Instrumente) was used to move the objective (Nikon 16×, 0.8 NA water immersion) in steps of 15 μm between frames to acquire images at 4 different depths. This resulted in an effective frame rate of 15 Hz. The field of view was approximately 375 by 300 μm. To chronically image the same site over days we marked recording locations in reference to landmarks based on the surface pattern of blood vessels.

For functional mapping of influence experiments, we performed simultaneous two-photon imaging and optogenetic stimulation of axons. The illumination source for stimulation was a 637 nm laser (Coherent, OBIS 637LX). Light was delivered to the sample in the same optical path as the excitation light through a dichroic mirror (ZT775sp-2p, Chroma). A long-pass dichroic mirror (F38-555SG, Semrock) was used to separate green emission light from stimulation and infrared excitation light. Light leak from the stimulation laser was further reduced by synchronizing the laser light output to the turnaround times of the resonant scanner (during which imaging data were not acquired). In combination with the 80 MHz digital band-pass filter, this allowed for stimulation-artifact-free synchronous imaging and optogenetic stimulation. Stimulation light (mean intensity of 6 or 12 mW on the sample) was pulsed at a frequency of 20 Hz with a 50% duty cycle. Stimuli were delivered for 1 s with an inter-trial interval of between 12 and 18 s during a closed loop and an open loop session.

### Two-photon data analysis

Data from each recording session were motion corrected for rigid X and Y displacement using cross-correlation. Briefly, we computed the cross-correlation between each frame and a reference template in the frequency domain to speed up computation and enable spatial frequency band-passing. The latter prevents vignetting, which impairs registration. Peak cross-correlation values were used to shift each frame. To improve robustness, the reference template was periodically updated by averaging the most recently aligned frames and image boundaries were cropped by 10% to reduce edge artifacts during registration. Regions of interest (ROIs) for each neuron were semi-automatically extracted from the mean and the maximum projection of the recording using a combination of Cellpose-based segmentation (Stringer et al. 2021) and custom-written software. For chronic recordings, ROIs were defined on the first recording day, semi-automatically matched across all time points and visually inspected. All ROIs that were not identifiable in all time points were discarded. Fluorescence traces were calculated as the mean pixel value of the ROI for each frame. Slow drifts were removed from the traces by high-pass filtering (Dombeck et al. 2007) with an intensity-preserving algorithm. A filtered trace was generated using the 8^th^ percentile computed in a moving window of 67 s (1000 frames). This filtered trace was median subtracted and then subtracted from the raw trace to generate a drift-corrected version. Activity was calculated as the ΔF/F_0_, where F_0_ was the median fluorescence over the entire trace.

Response onset plots for grating stimuli and optogenetic stimulation (**Figure 5B, 6C, 6E, 7B-D**) were baseline subtracted in a window from -0.5 to 0 s. To calculate reliability for a given stimulus, we separately averaged neuronal response to all even and odd repetitions of that stimulus and then computed the correlation coefficient between the two baseline-subtracted traces in a window of -1 to 3 s (relative to stimulus onset) for gratings and -1 to 2 s for optogenetic stimulation. Response amplitude was calculated as the mean of the baseline-subtracted response to all events in a window of 0.5 to 2 s for grating and optogenetic stimulation experiments. In a subset of plots (**Figure 5D, 6G, S5A-B, S8A-C**), amplitudes were then transformed using a symmetric logarithmic function,

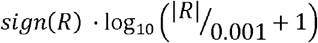

to visualize both positive and negative values across a wide dynamic range while preserving linearity near zero.

To quantify orientation selectivity (**Figure 7F-G**), we fitted a circular normal tuning curve model to each neuron’s responses across the 8 grating directions. The model is defined as:

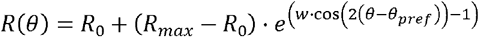

*R*_*0*_ is the baseline response, *R*_max_ is the peak response at the preferred orientation, *θ* _*pref*_ is the preferred orientation, and *w* is the concentration parameter controlling tuning width. Model parameters were estimated by minimizing the residual sum of squares. To determine whether a neuron was significantly orientation selective, we compared the fitted model to a null model consisting of a flat tuning curve at the mean response amplitude. For each model, we computed the penalized likelihood criterion (PLC):

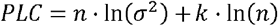

where *n* is the number of data points, *σ* ^2^ is the residual variance, and *k* is the number of free parameters (1 for the null model, 4 for the circular normal model). We then calculated *ΔPLC = PLC*_*circ norm*_ *− PLC*_*null*_, where positive values indicate that the circular normal model provides a better fit. To establish a significance threshold, we generated a null distribution for each neuron by randomly permuting the direction labels and refitting the model 500 times. A neuron was classified as orientation-selective if its actual measured ΔPLC exceeded the 95^th^ percentile of its own shuffle distribution. Neurons with *w* ≤ ln(2) were excluded from tuning width analyses, as this corresponds to tuning curves broader than 180° full width at half maximum, indicating negligible orientation selectivity. Tuning width was quantified as the full width at half maximum of the fitted tuning curve, derived analytically from the concentration parameter *w. For the a*nalysis in Figure 7G, mice with fewer than 5 orientation-selective neurons were not included.

### Widefield calcium imaging

Widefield imaging experiments were performed on a custom-built macroscope consisting of commercially available objectives mounted face-to-face (Nikon 85 mm f/1.8 sample side, Nikon 50 mm f/1.4 sensor side). We used a 470 nm LED (Thorlabs), powered by a custom-built LED driver for exciting GCaMP fluorescence through an excitation filter (SP490, Thorlabs) reflected off of a dichroic mirror (LP490, Thorlabs) placed in the parfocal plane of the objectives. Green fluorescence was collected through a 525/50 nm emission filter on an sCMOS camera (PCO edge 4.2, Excelitas Technologies Corp). Apertures on objectives were usually kept maximally open and the current at the LED driver was used to adjust fluorescence intensity to a value that was kept below 25% of the maximum dynamic range of the sensor. LED illumination was adjusted with a collimator (Thorlabs SM2F32-A) to achieve homogeneous illumination across the surface of the cranial window. The resulting normal profile of the illumination cone was further trimmed with black tape on the sample side objective to avoid directly illuminating regions outside of the craniotomy. An Arduino board (Arduino Mega 2560) was used to control LED onsets to the frame trigger signal of the camera. The duty cycle of the 470 nm LED was 90%. Raw images were acquired at 100 Hz and full dynamic range (16 bit) of the sensor. Raw images were cropped on-sensor and the resulting data were streamed to disk with custom software written in LabVIEW, resulting in an effective pixel size of 60 μm^2^ at a standardized imaging resolution of 1108 by 1220 pixels (1.35 MP).

### Widefield imaging data analysis

For analysis of chronic widefield macroscope imaging, we first aligned raw movie data across time points. We used MATLAB’s imregtform function to calculate a transformation matrix from a single frame relative to a reference frame from the first imaging session. Because no detectable movement occurred within imaging sessions, we applied this transformation matrix to the entire raw image sequence. We used a similarity threshold of 0.5 as a pass criterion, followed by visual inspection. In rare cases where the alignment algorithm did not converge above threshold, or visual inspection revealed misalignment, we manually adjusted the starting points for imregtform until satisfactory alignment was achieved. The portion of the images containing dorsal cortex was then tiled with 818 ROIs each with a fixed location relative to user-defined bregma and lambda. This resulted in an average ROI size of 25 by 25 pixels (approximately 190 by 190 μm). Each ROI was also registered against the Allen Mouse Brain Common Coordinate Framework (CCFv3-2017) (Wang et al. 2020). Fluorescence traces for each ROI were calculated by averaging all pixel values. Slow drifts were removed from the traces by high-pass filtering (Dombeck et al. 2007) with an intensity-preserving algorithm. A filtered trace was generated using the 8^th^ percentile computed in a moving window of 62.5 s (6250 frames). This filtered trace was median subtracted and then subtracted from the raw trace to generate a drift-corrected version. Activity was calculated as the ΔF/F_0_, where F_0_ was the median fluorescence of the entire trace.

To analyze the activity correlation structure (**Figure 2B, 2C, 3F, 3G**), we used data from all available virtual reality paradigms: closed loop, open loop, dark, and gratings. We first calculated the correlation coefficient of neuronal activity between all possible pairs of ROIs. We then determined the distance between each pair of ROIs and normalized this value by the distance between bregma and lambda obtained from the images during initial surgery for each mouse. We plotted the correlation against the distance for each pair of ROIs and binned the data in a 40 by 40 grid and smoothed the resulting image with a Gaussian filter (size: 11 by 11, sigma: 2.5). Contour lines in the heatmaps were then drawn to 50% of the maximum value in the naive plot. The change in correlation was calculated for each ROI pair as the correlation at a given time point minus baseline, where baseline is the grand mean of the correlation coefficients for that ROI pair across all naive sessions and mice.

### Histology

Mice were deeply anesthetized and perfused via intracardiac infusion with room temperature PBS and then 4% paraformaldehyde (PFA, w/vol, dissolved in 0.1 M phosphate buffer, pH 7.4). Brains were extracted and post-fixed overnight in PFA 4% at 4 C, then stored in PBS at 4 C. Brains were then embedded in 4% low melting agarose (Biozym, 850081) and coronal sections (50-60 μm thick) of the brain region of interest were cut using a vibratome (Leica VT1000 S) and collected in a multi-well plate. Sections were mounted on microscopy slides with a mounting medium (VECTASHIELD Plus antifade mounting medium, H-1900, Vector Laboratories) and stored at 4 C. Slides were imaged on an upright microscope (Axio Imager M2, Zeiss) equipped with a confocal scanner (Yokogawa) with a 50 μm pinhole. Illumination was achieved with a 561 nm (Cobolt) and a 488 nm (Toptica Photonics) laser. A plan apochromat 20×/0.8NA objective (Zeiss) was used, resulting in an effective pixel size of 0.325 μm. Up to 3 optical sections with an interval of 15 μm were acquired with a Z-piezo (Applied Scientific Instrumentation). Neuron positions in the histological images were estimated as previously described (Cicconet and Hochbaum 2019; Hrvatin et al. 2020). Briefly, we used Laplacian-of-Gaussian filtering to detect local maxima candidates, followed by correlation-based selection against an ideal 2D Gaussian. The boundary for the cortex surface was automatically detected based on intensity and manually inspected for each image. Cortical depth of a neuron was defined as the minimum distance between the neuron’s position and the cortex boundary.

Mice injected in thalamus were classified into predominantly first-order or higher-order thalamus labeling groups based on visual inspection of histological images, performed blind to functional data. To validate this classification, we quantified mean fluorescence intensity within a region of interest in the dLGN for a subset of mice, confirming that the two groups differed in the degree of viral labeling in first-order visual thalamus. Mice in which the injection site was mislocalized outside thalamus were excluded from the analysis.

### Slice preparation for patch-clamp recordings

Slice electrophysiology experiments were performed in Tlx3-Cre x Ai14 mice aged between 45 and 55 d. Between 10 and 14 days prior to patch-clamp experiments, we performed intracortical injections of an AAV to express EGFP in a Cre-dependent manner. Mice were first anesthetized with isoflurane (4% in O_2_, Vapor, Draeger) and then immediately decapitated. To increase cell viability, mice were exposed to an oxygen-enriched atmosphere for at least 10 min before decapitation. The brain was dissected in ice-cold sucrose-based solution (at about 1 °C), and coronal brain slices (300 μm thick) were cut using a vibratome. For cutting and storage, a sucrose-based solution was used, containing (in mM): 87 NaCl, 25 NaHCO_3_, 2.5 KCl, 1.25 NaH_2_PO_4_, 75 sucrose, 0.5 CaCl_2_, 7 MgCl2, and 10 glucose (equilibrated with 95% O_2_ / 5% CO_2_, osmolarity 340 mOsm). Slices were kept in a holding chamber at 35 °C for 30 min after slicing and subsequently stored at room temperature until experiments were performed.

### Whole-cell recordings

Electrophysiological recordings of acute brain slices were conducted within about 6 hours after slice preparation. Slices were transferred into a recording chamber and continuously perfused with oxygenated ACSF. The ACSF was maintained at approximately 32 °C and contained (in mM): 125 NaCl, 25 NaHCO_3_, 25 glucose, 2.5 KCl, 1.25 NaH_2_PO_4_, 2 CaCl_2_, and 1 MgCl_2_ (equilibrated with 95% O_2_/5% CO_2_, osmolarity 320 mOsm). L5 pyramidal cells were visually identified in L5 of V1 using infrared differential interference contrast video microscopy and a 40× objective. Developmentally and adult labeled Tlx3 cells were identified using red (tdTomato) and green (EGFP) fluorescence using a cooled CCD camera system (SensiCam, TILL Photonics, Gräfelfing, Germany). Excitation light was provided by a source with a wavelength of 555 and 470 nm (Polychrome V, TILL Photonics) connected to the microscope using a quartz optic fiber light guide. The illumination intensity was kept low to avoid possible photo-bleaching of the neurons and subsequent phototoxicity. Voltage- and current-clamp recordings were conducted with patch pipettes (4-6 MΩ) pulled from borosilicate glass tubes with 2.0 mm outer diameter and 0.5 mm wall thickness (Hilgenberg, Malsfeld, Germany; Flaming-Brown P-97 puller, Sutter Instruments, Novato, USA). For current-clamp recordings of the intrinsic properties, patch pipettes were filled with a potassium-based intracellular solution containing (in mM): 160 K-gluconate, 0.1 EGTA, 10 HEPES, 2 MgCl_2_, 2 Na_2_ATP, 0.3 GTP, 1 phosphocreatine and 0.5% biocytin, adjusted to pH 7.3 with KOH. These experiments were performed in the presence of ionotropic receptor blockers 10 μM NBQX, 25 μM D-AP5, and 10 μM gabazine. Voltage- and current-clamp recordings were both obtained using a Multiclamp 700B amplifier (Molecular Devices, Palo Alto, CA, USA), filtered at 10 kHz, and digitized at 20 kHz with a CED Power 1401 interface (Cambridge Electronic Design, Cambridge, UK). Data acquisition was controlled using IGOR Pro 6.31 (WaveMetrics, Lake Oswego, Oregon) and the CFS library support from CED (Cambridge Electronic Design, Cambridge, UK). Stock solutions of NBQX (Tocris Bioscience, 10 mM), D-AP5 (Tocris Bioscience, 25 mM) and gabazine (SR 95531 hydrobromide, Sigma-Aldrich, 10 mM) were dissolved in water. All drugs were stored as aliquots at -20 °C and diluted in ACSF.

Series resistance (R_series_) was determined by giving a -5 mV voltage step for 500 ms in voltage-clamp mode (command potential set at -70 mV). R_series_ was calculated by dividing the -5 mV voltage step by the peak current value generated immediately after the step in the command potential. Input resistance (R_input_) was measured in current-clamp mode by applying a -25 pA hyperpolarizing current step and calculating the resulting steady-state voltage change according to 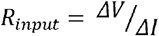 I where ΔV was defined as the difference between the baseline membrane potential (command potential set at -70 mV) and the steady-state voltage at the end of the current pulse.

The resting membrane potential was calculated by holding the neuron in voltage-follower mode (current-clamp, at I = 0) immediately after breaking in and averaging the membrane potential over the next 20 s. Depolarizing current pulses (25 pA increments of 1 s duration) in current-clamp were injected to examine each neuron’s basic suprathreshold electrophysiological properties (baseline membrane potential was maintained at -70 mV, by injecting the required current, when necessary). Action potential width was calculated as the full width at the half-maximum amplitude of the first resulting action potential at rheobase. The action potential threshold was measured as the membrane potential at which the depolarization slope exhibited the first abrupt change (Δslope > 10 V/s). Current threshold was defined as the minimum current injection for action potential generation. Membrane time constant was calculated by fitting an exponential function to the membrane potential trace after applying a 25 pA negative current pulse at rest (-70 mV). Membrane capacitance (C_m_) was measured in whole-cell voltage-clamp configuration using the capacitive transient evoked by a small voltage step. Cells were held at the constant holding potential of −70 mV, and brief voltage steps of −5 mV were applied to minimize activation of voltage-dependent conductances. The resulting capacitive transient currents were recorded and leak-subtracted. Total charge transfer (Q) associated with the capacitive transient was calculated by integrating the transient current over time. Membrane capacitance was then calculated according to the relation 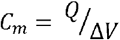, where ΔV is the amplitude of the voltage step. Adaptation ratio of action potentials was calculated at the current step with the maximal action potential firing by dividing the average interspike interval (ISI) of the last 5 ISIs by the 1^st^ ISI. For the analysis of the number of action potentials as a function of current, cells that had fewer than 5 total action potentials over the course of the entire experiment were excluded.

Data analysis for patch-clamp data was performed offline using the open-source analysis software Stimfit and custom scripts written in Python.

### Generation of single cell suspensions for RNA sequencing

We generated single cell suspensions as previously described (O’Toole et al. 2023), adapted from (Hempel et al. 2007; Tasic et al. 2018). Throughout the experiment we used N-Methyl-D-glucamine (NMDG)-based ACSF to improve cell viability (Ting et al. 2014). The NMDG ACSF formulation consisted of NMDG (96 mM), NaHCO3 (25 mM), HEPES (20 mM), N-acetylcysteine (12 mM), MgSO4 (10 mM), glucose (25 mM), sodium L-ascorbate (5 mM), myo-inositol (3 mM), sodium pyruvate (3 mM), KCl (2.5 mM), thiourea (2 mM), NaH2PO4 (1.25 mM), CaCl2 (0.5 mM), and taurine (0.01 mM). All ACSF stocks were titrated with HCl (5 M) to pH 7.35 and kept at 315 mOsm. Additionally, all solutions were supplemented with 13.2 mM trehalose to enhance cell survival during the dissociation and sorting (Saxena et al. 2012). Solutions were supplemented with a transcriptional blocker, actinomycin-D (1 μg/mL), to reduce dissociation-related expression changes (Wu et al. 2017). In addition, the ion channel blockers TTX (0.1 μM), DNQX (20 μM), and APV (50 μM) were added to prevent any activity-related changes and to reduce excitotoxicity (Hempel et al. 2007). The ACSF used for slicing and cooling the brain was pre-chilled to an icy slush and bubbled with carbogen gas (95% O_2_, 5% CO_2_) throughout the experiment.

To generate single cell suspensions, mice were anesthetized with isoflurane and decapitated. We removed the brain, glued it to a cutting platform next to an agarose block and rapidly immersed it in ice-cold carbogenated ACSF. Brain sections (300 μm) were cut on a vibratome. Sections within and around V1 were collected with a plastic transfer pipette and placed into a bubbled chamber filled with room temperature ACSF and allowed to recover for at least 5 min. Subsequently, samples were placed into ACSF with 1 mg/mL Protease Type XIV and 33.3 μg/mL collagenase, and digested for 60 min. Slices were placed in ACSF (without protease/collagenase) and allowed to recover for 15 min. After that, the slices were transferred to a Petri dish containing cold carbogenated ACSF supplemented with 1% Fetal Bovine Serum (FBS). For each slice, a portion of V1 L5 containing fluorescent EGFP labeling was microdissected under an epifluorescence microscope. Care was taken to microdissect only parts of the cortical tissue that exhibited EGFP fluorescence. Microdissected pieces were then transferred to 1.5 Ml RNase-free tubes containing ACSF supplemented with 1% FBS. These tissue pieces were triturated with a series of fire-polished glass transfer pipettes, with inner diameters of 600 μm, 300 μm, and 150 μm. Single-cell suspensions were then strained through nylon filters (Celltrics filters, Sysmex) and placed into tubes along with 2 μL of a live/dead cell stain (DRAQ7, BioLegend) before FACS sorting.

### Flow cytometry

Cells were analyzed and sorted using a BD FACSAria III controlled by BD FACSDiva software (version 8.0.1). A sequential gating strategy was applied to identify live, single cells. First, intact cells were selected based on forward scatter area and side scatter area to exclude debris. Doublets were then excluded using forward scatter height versus width and side scatter height versus width. Finally, live cells were identified by exclusion of DRAQ7, a membrane-impermeant DNA dye that labels cells with compromised membranes. The resulting population was sorted based on red and green fluorescence. Gate boundaries were established in a pilot experiment and applied consistently across all samples.

### RNA sequencing

RNA for bulk RNA-sequencing was purified with the Single Cell RNA Purification Kit (Norgen), and columns were treated with a DNase to remove genomic DNA (Norgen). The isolation was performed according to the manufacturer’s instructions. Purified RNA was assessed for integrity using Bioanalyzer Pico RNA chips (Agilent). Purified RNA was used as input for full-length cDNA amplification using Smart-seq2 (oligo-dT priming and template-switching, as described in (Picelli, Faridani, et al. 2014), with minor modifications). Purified cDNA (1 ng) was tagmented using in-house purified Tn5 transposase (Picelli, Björklund, et al. 2014) in TAPS-DMF buffer and amplified using Illumina Unique Dual Index primers. Libraries were sequenced on an Illumina NovaSeq6000 using 112 cycles, paired-end (2 × 56 bp). Demultiplexing was performed using bcl2fastq2 v2.20.

### RNA-sequencing analysis

Initial RNA-sequencing analysis was conducted on a local Galaxy server (version 21.05) (The Galaxy Community 2024) executing several R scripts (R version 4.4.1 and Bioconductor version 3.19): paired-end sequence reads were aligned to the mouse genome (GRCm39 primary assembly) using the “qAlign” function from the Bioconductor package QuasR version 1.44.0 (Gaidatzis et al. 2015) with default parameters except for aligner = “Rhisat2”, splicedAlignment=“TRUE”, allowing only uniquely mapping reads. Raw gene counts were generated using the “qCount” function (QuasR) with GENCODE release M28 ‘basic gene annotation’ (“gencode.vM28.annotation.gtf” as TxDb object) as query, with default parameters except useRead=first and orientation=any. The count table was modified by replacing the Ensembl gene ID with the gene symbol (if unique) or a combination of the gene symbol and the Ensembl ID (if the gene symbol was not unique) or keeping the Ensembl ID (for genes without a gene symbol) as the unique identifier. The count table was filtered to only keep genes which had at least 0.5 counts per million (CPM) in at least 5 samples. Differential gene expression was calculated with the Bioconductor package edgeR (version 4.2.1, (Lun et al. 2016)) using the quasi-likelihood F-test after applying the calcNormFactors function, obtaining the dispersion estimates and fitting the negative binomial generalized linear models between the two groups.

Cell type deconvolution of the bulk RNA-sequencing data was performed with the R package BayesPrism (version: 2.2.2) (Chu et al. 2022) according to the vignette (R version 4.5.2 and Bioconductor version 3.22). The “Mouse Whole Cortex and Hippocampus 10x” dataset from the Allen Institute (Yao et al. 2021) was used as the scRNA-sequencing reference. We used a grouping level defined in the reference dataset which aggregates cell subclasses in 8 different groups named neighborhoods. Prior to filtering and marker gene selection, the single cell reference was down-sampled to include only 3000 cells for each of the 8 neighborhood labels. The prism object was built using the neighborhood labels as “cell.type.labels” and “cell.state.labels” with “key=NULL”, “outlier.cut=0.01”, and “outlier.fraction=0.1”. The deconvolved cell type fractions were extracted from the output of “run.prism” with which.theta=“final”, state.or.type=“type”.

### Statistical analysis

All quantitative information for the statistical analyses performed in this manuscript is provided in **Table S1**. Sample sizes for all experiments were determined based on effect sizes observed in preliminary data or reported in the published literature. Blinding of experimenters to the treatment condition was not feasible due to the behavioral sedation produced by the administered compounds. We applied a uniform baseline-matching procedure across all experiments: Using exclusively data from the naive time point, and therefore blind to treatment outcomes, we iteratively excluded mice with the most extreme baseline values, alternating between groups, until the two distributions were statistically indistinguishable. Across the entire study, this procedure resulted in the exclusion of only a single mouse in Figure 5, as all other experiments required no exclusions. For all statistical tests involving hierarchically structured data we used a hierarchical bootstrapping approach to account for dependencies between groups of data points (Saravanan et al. 2020). In these cases, the data were first resampled with replacement at the level of mouse or recording site and then resampled with replacement at the level of ROIs or neurons. We performed 10 000 bootstrap repetitions and calculated a distribution of bootstrapped mean values. The p value was calculated as the fraction of the bootstrap sample consistent with the null hypothesis. All statistical testing for non-hierarchical datasets was performed as follows. To test if distributions are significantly different from zero, or if two paired distributions are different from each other, we used a signed rank test when sample sizes were sufficient (n ≥ 6 allowing detection at α = 0.05). For paired data and smaller sample sizes, we used paired t-tests (Winter 2013). To test if two unpaired distributions are different from each other, we used a rank-sum test. To analyze non-hierarchical paired datasets with more than one experimental variable we used a two-way repeated measures ANOVA followed by post-hoc testing adjusted for false discovery rate (FDR) as previously described (Benjamini and Hochberg 1995). In cases where one of the two factors had incomplete data due to experimental constraints (i.e., sedation of mice; **Figure S1**), we used a linear mixed-effects model to accommodate the unbalanced design.

## Supplementary figures

**Figure S1.**
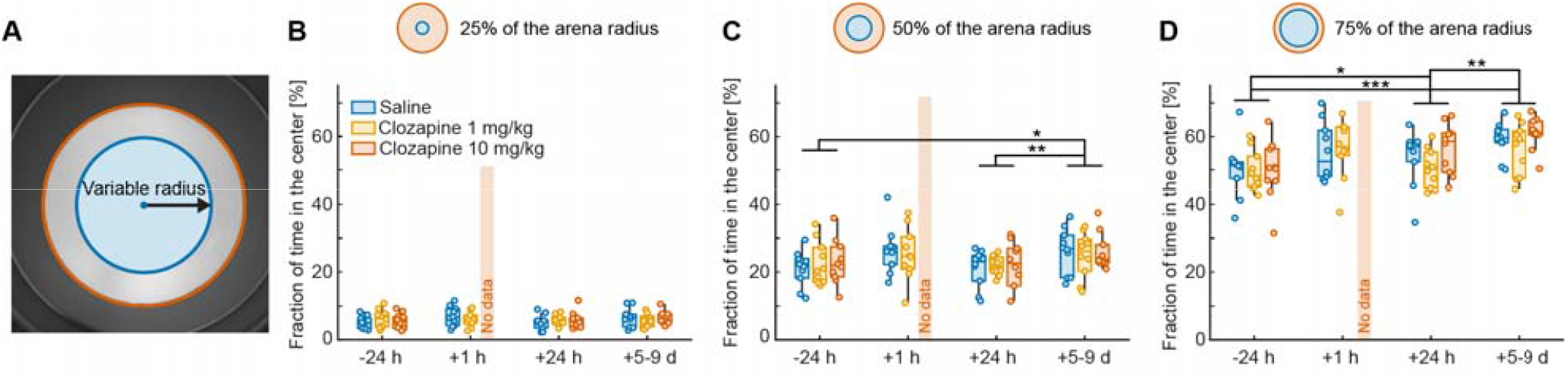
Clozapine does not alter thigmotaxis in mice. **(A)**For analysis, the arena (orange circle) was divided into center (blue area) and periphery. Quantification was performed as a function of the relative radius of the center. **(B)**Fraction of time spent in the center of the arena (25% of arena radius) per session. Data from mice injected with clozapine (10 mg/kg) at the +1 h time point were excluded due to sedation (**Figure 1B, C**); these mice exhibited minimal locomotion and remained largely stationary at their initial placement, precluding meaningful interpretation of center-zone occupancy. *: p < 0.05, **: p < 0.01, ***: p < 0.001. See **Table S1** for all statistical information. **(C)**As in **B**, but the center is a circle with a radius that is 50% of the arena. **(D)** As in **B**, but the center is a circle with a radius that is 75% of the arena.

**Figure S2.**
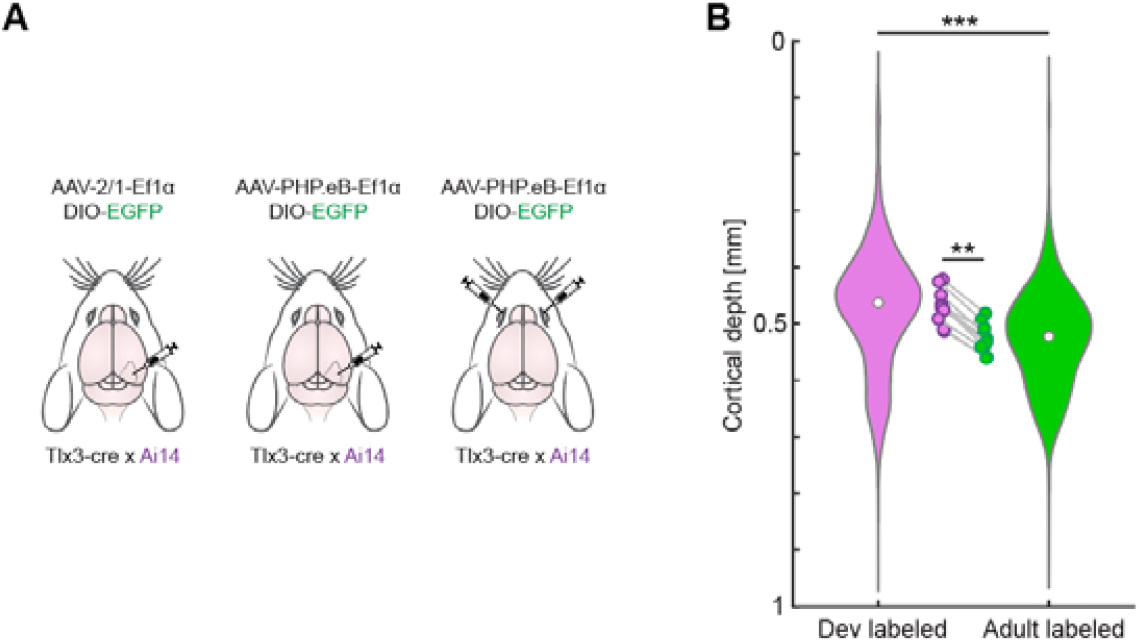
Depth distribution of all developmentally and adult labeled Tlx3 neurons. **(A)** Schematic of the experiment. **(B)** Depth distribution of developmentally and adult labeled Tlx3 neurons in V1 aggregated across three viral labeling strategies, violins represent the distribution of depth for all neurons, dots the median of the distributions, and data points each mouse’s average. *: p < 0.05, **: p < 0.01, ***: p < 0.001. See **Table S1** for all statistical information.

**Figure S3.**
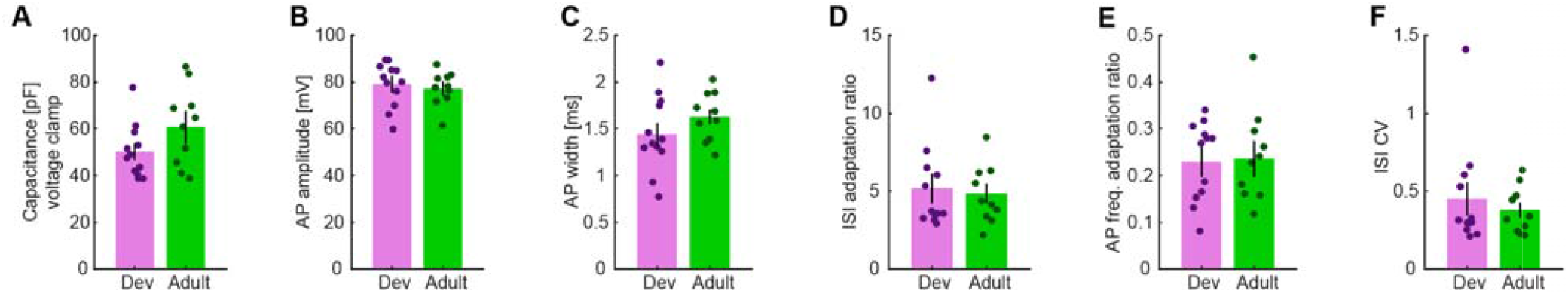
Similar intrinsic physiological properties between developmentally and adult labeled Tlx3 neurons. **(A)** Membrane capacitance in voltage-clamp configuration in developmentally and adult labeled Tlx3 neurons. Here and elsewhere bar height represents the mean of the distribution of bootstrapped means, error bars the standard deviation of the distribution of bootstrapped means, data points individual cells. **(B)** As in **A**, but showing amplitude of action potentials. (**C**) As in **A**, but showing the width of action potentials. **(D)** As in **A**, but showing the adaptation ratio of the interspike interval. Differences in the adaptation ratio reflect different bursting profiles of the neurons. **(E)** As in **A**, but showing the adaptation ratio of the action potential frequency. **(F)** As in **A**, but showing the coefficient of variation for the interspike interval.

**Figure S4.**
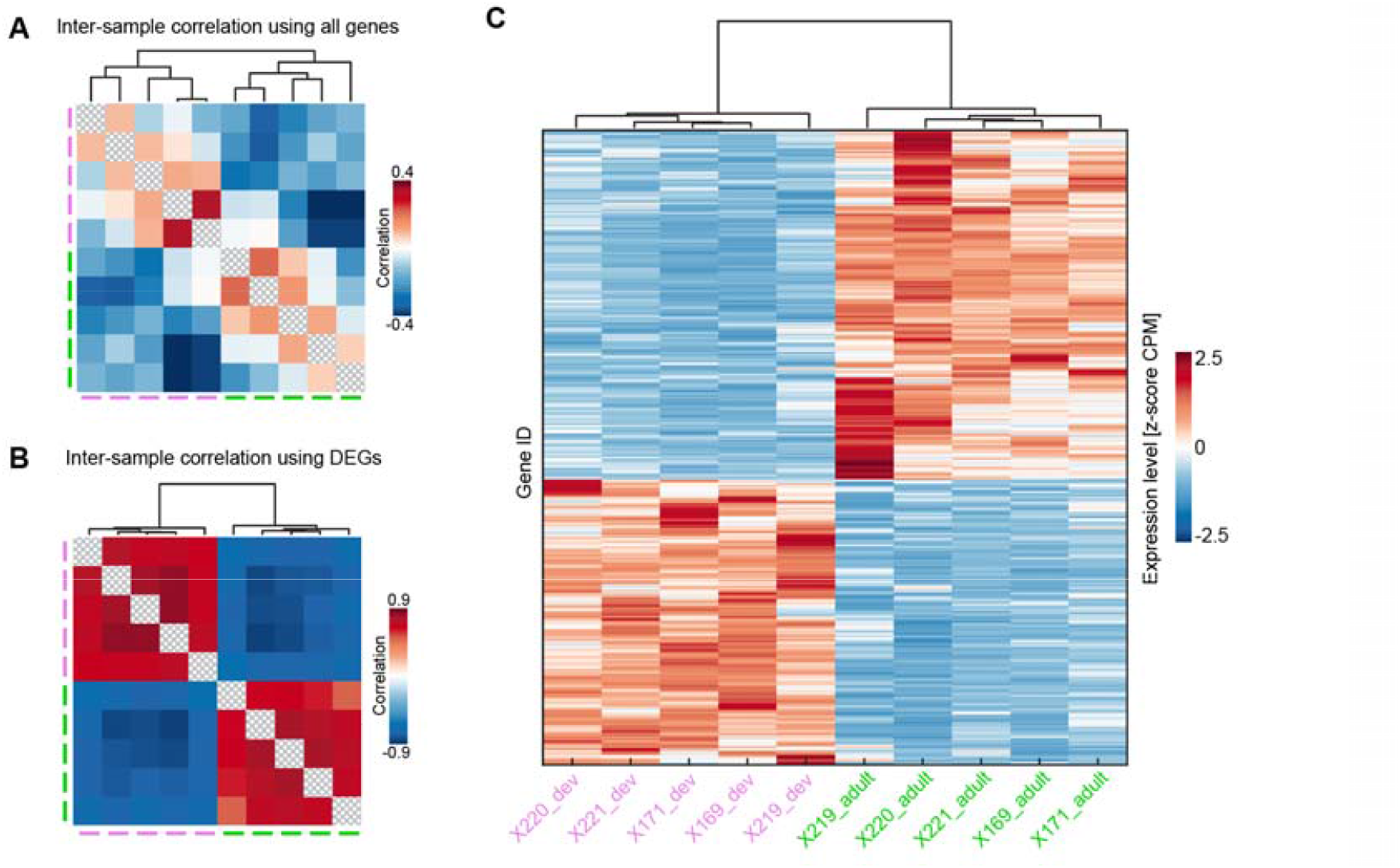
The set of differentially expressed genes was similar across samples. **(A)**Sample-to-sample correlation based on all filtered (≥ 0.5 CPM in at least 5 samples) genes (17 682 genes). Correlation coefficients were computed on z-scored expression values (CPM). Color labels for columns and rows indicate whether the sample was from developmentally (magenta) or adult labeled (green) Tlx3 neurons. Data in the main diagonal were excluded from the plot. Here and elsewhere the dendrogram shows hierarchical clustering of samples based on expression profiles (average linkage, correlation distance). **(B)**As in **A**, but showing sample-to-sample correlation based on differentially expressed genes (FDR < 0.05, |log_2_FC| > 1, 445 genes). **(C)**Heatmap of differentially expressed genes across individual samples. Expression values (CPM) were z-scored by gene. Columns represent samples from developmentally (dev, magenta) or adult labeled (adult, green) Tlx3 neurons.

**Figure S5.**
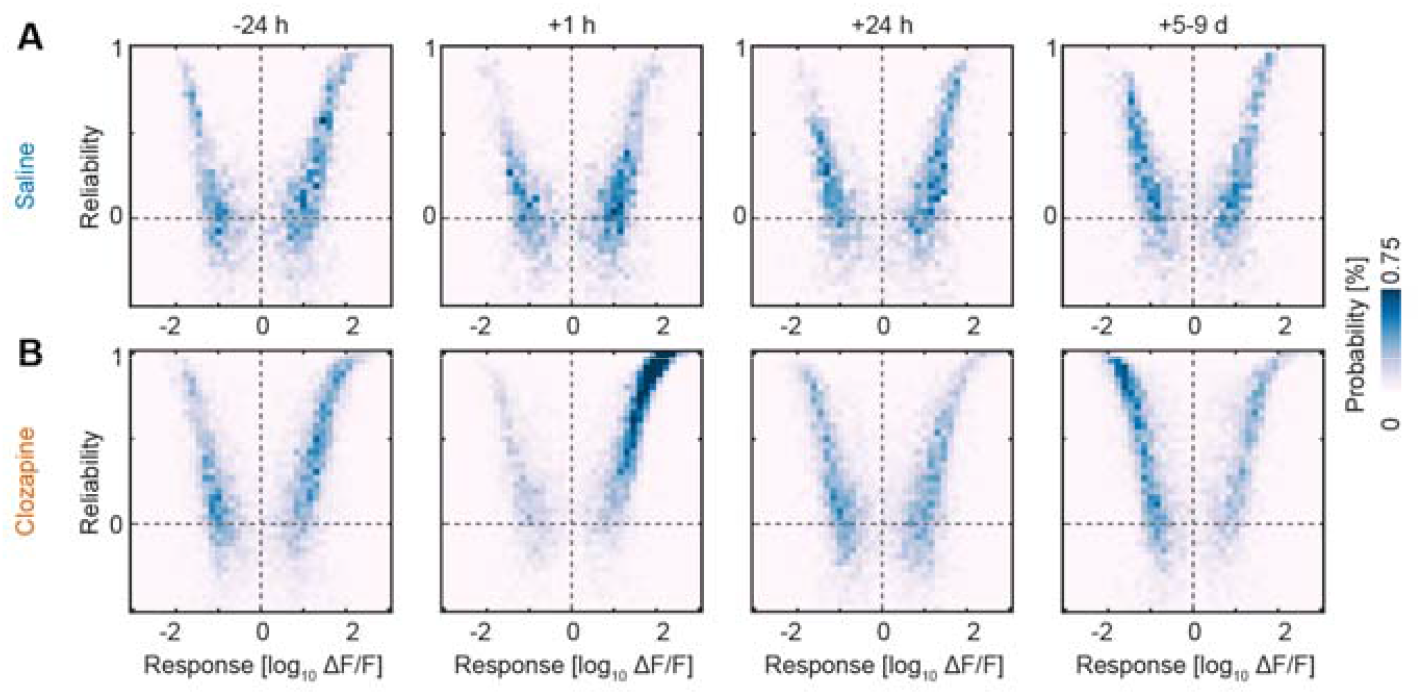
Distributions of reliability and response amplitude to optogenetic stimulation of cingulate axons over time. **(A)**Density plot (40 by 40 bins) showing the distribution of V1 neurons from saline injected mice in the space of reliability and response amplitude to the optogenetic stimulation of cingulate axons for different time points after injection. This illustrates the underlying distributions used to compute the difference in **Figure 5D**. **(B)**As in **A**, but for clozapine injected mice.

**Figure S6.**
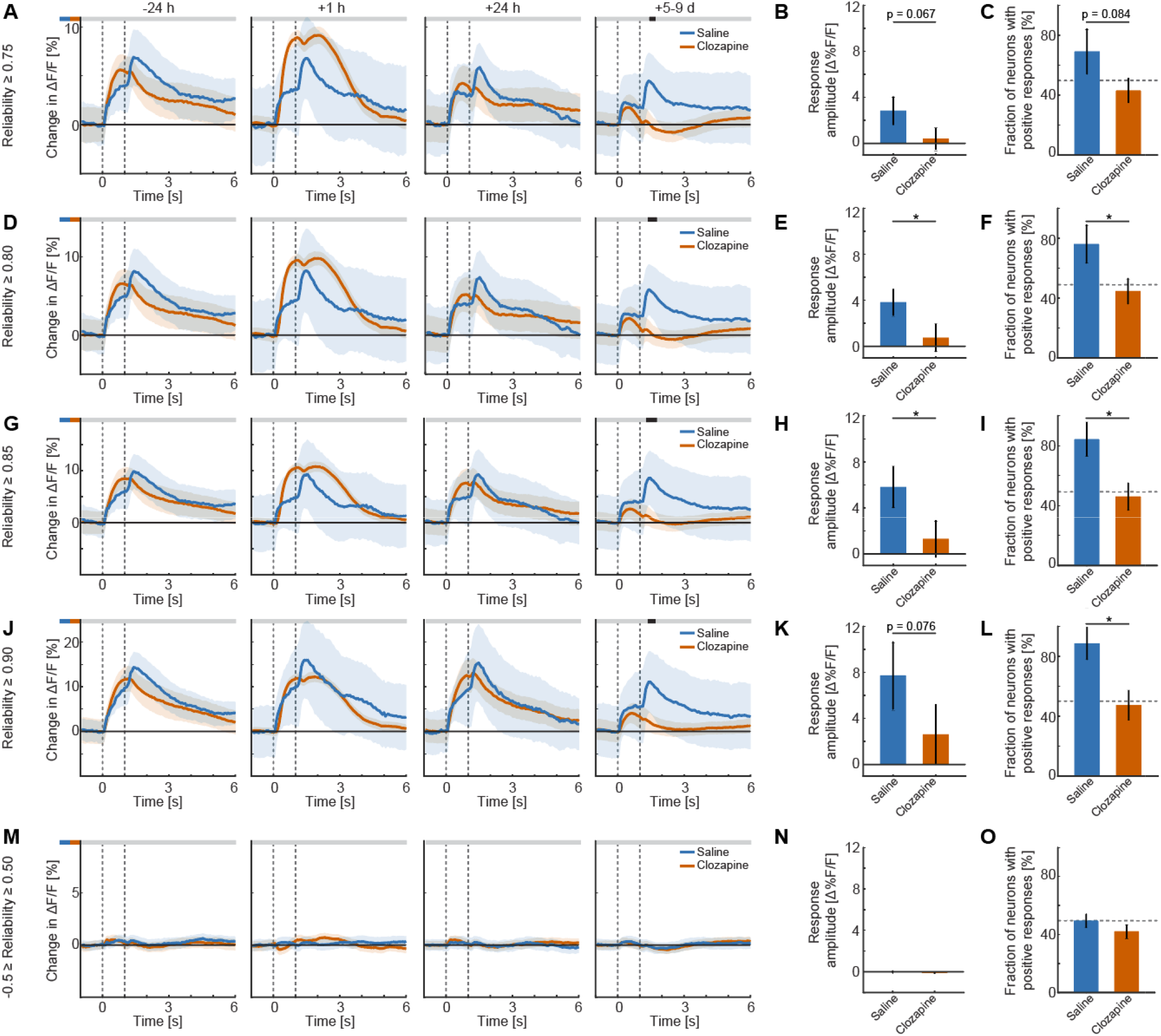
The long-term effect of clozapine is robust to different reliability thresholds. **(A)**Population responses to optogenetic stimulation of neurons with reliability ≥ 0.75, measured before and at various time points after injection. Shading indicates ± 1 standard deviation of the hierarchically bootstrapped distribution. Vertical dashed lines mark stimulus onset and offset. The horizontal bar above the plot indicates time bins with statistically significant differences between saline and clozapine injected mice (gray: not significant; black: p < 0.05). See **Table S1** for all statistical information. **(B)**Mean response amplitude to optogenetic stimulation 5-9 days after injection for neurons with reliability ≥ 0.75. Bar height and error bars represent the mean and standard deviation of the bootstrapped mean distribution, respectively. *: p < 0.05, **: p < 0.01, ***: p < 0.001. **(C)**Fraction of neurons with reliability ≥ 0.75 showing positive responses 5-9 days after injection. Bar height and error bars represent the mean and standard deviation of the bootstrapped mean distribution, respectively. **(D)**As in **A**, but for neurons with reliability ≥ 0.80. **(E)** As in **B**, but for neurons with reliability ≥ 0.80. **(F)** As in **C**, but for neurons with reliability ≥ 0.80. **(G)** As in **A**, but for neurons with reliability ≥ 0.85. **(H)** As in **B**, but for neurons with reliability ≥ 0.85. **(I)** As in **C**, but for neurons with reliability ≥ 0.85. **(J)** As in **A**, but for neurons with reliability ≥ 0.90. **(K)** As in **B**, but for neurons with reliability ≥ 0.90. **(L)** As in **C**, but for neurons with reliability ≥ 0.90. **(M)** As in **A**, but for neurons with reliability between -0.5 and 0.5. **(N)** As in **B**, but for neurons with reliability between -0.5 and 0.5. **(O)** As in **C**, but for neurons with reliability between -0.5 and 0.5.

**Figure S7.**
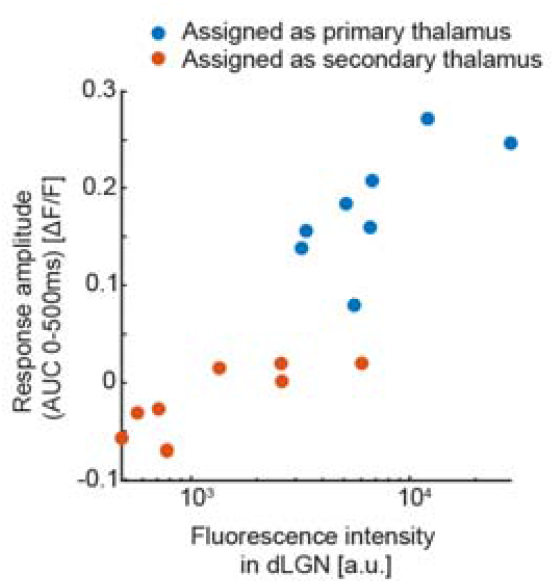
Functional responses depend on whether injection preferentially occurred in first-order or higher-order thalamus. Scatter plot showing the average early response amplitude (0-500 ms) and mean red (ChrimsonR) fluorescence intensity in the dorsal lateral geniculate nucleus for each mouse. Color represents the injection group that each mouse was assigned to in **Figure 6**.

**Figure S8.**
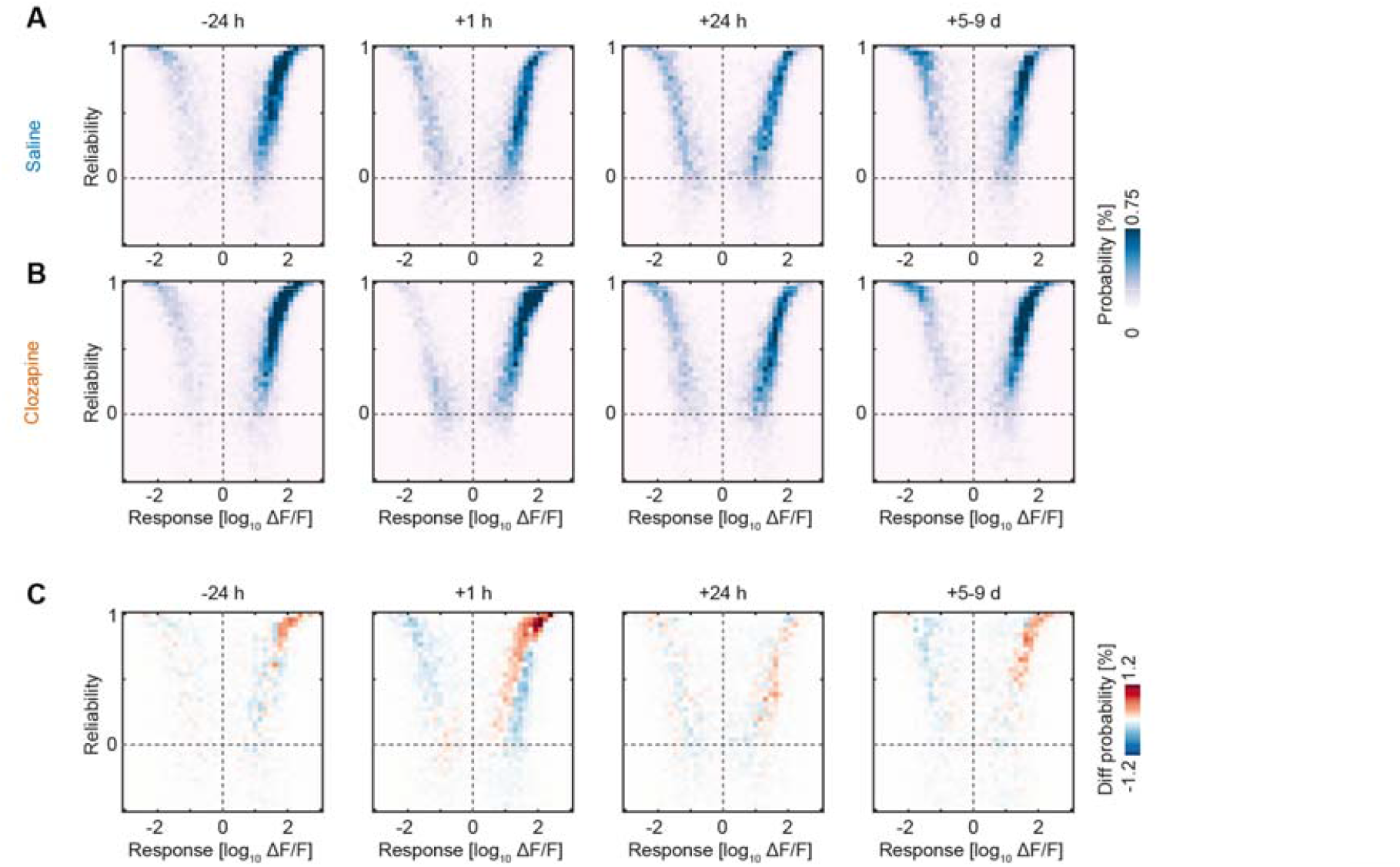
Reliability heatmaps of the responses to gratings. **(A)**Density plot (40 by 40 bins) showing the distribution of V1 neurons from saline injected mice in the space of reliability and response amplitude to full field gratings for different time points after injection. **(B)**As in **A**, but for clozapine injected mice. **(C)**As in **A**, but showing the difference between distributions of saline and clozapine injected mice.

## Supplementary tables

**Table S1.** Statistical information. For a description of all the statistical tests used, please refer to the **Methods and Materials** section, paragraph **Statistical analysis**. False discovery rate (FDR) estimates were performed as previously described (Benjamini and Hochberg 1995). In case of FDR adjustment, the p value given is without adjustment, p_adj_ the one with.

| Figure | Hypothesis tested | Test | N (per group) | Unit | p-value |
|---|---|---|---|---|---|
| **1C** | Total distance traveled different between groups | Two-way repeated measures ANOVA | 10, 10, 10 | Mice | Time*treatment:<br>p < 0.001 |
|  | -24 h, saline vs clozapine (1 mg/kg) | Post-hoc pairwise, FDR adjusted | 10, 10 | Mice | p = 0.21<br>p<sub>adj</sub> = 0.28 |
|  | -24 h, saline vs clozapine (10 mg/kg) | Post-hoc pairwise, FDR adjusted | 10, 10 | Mice | p = 0.61<br>p<sub>adj</sub> = 0.61 |
|  | -24 h, clozapine (1 mg/kg) vs clozapine (10 mg/kg) | Post-hoc pairwise, FDR adjusted | 10, 10 | Mice | p = 0.45<br>p<sub>adj</sub> = 0.49 |
|  | +1 h, saline vs clozapine (1 mg/kg) | Post-hoc pairwise, FDR adjusted | 10, 10 | Mice | p < 0.001<br>p<sub>adj</sub> < 0.0015 |
|  | +1 h, saline vs clozapine (10 mg/kg) | Post-hoc pairwise, FDR adjusted | 10, 10 | Mice | p < 0.001<br>p<sub>adj</sub> < 0.001 |
|  | +1 h, clozapine (1 mg/kg) vs clozapine (10 mg/kg) | Post-hoc pairwise, FDR adjusted | 10, 10 | Mice | p < 0.001<br>p<sub>adj</sub> < 0.001 |
|  | +24 h, saline vs clozapine (1 mg/kg) | Post-hoc pairwise, FDR adjusted | 10, 10 | Mice | p = 0.12<br>p<sub>adj</sub> = 0.18 |
|  | +24 h, saline vs clozapine (10 mg/kg) | Post-hoc pairwise, FDR adjusted | 10, 10 | Mice | p < 0.019<br>p<sub>adj</sub> < 0.044 |
|  | +24 h, clozapine (1 mg/kg) vs clozapine (10 mg/kg) | Post-hoc pairwise, FDR adjusted | 10, 10 | Mice | p = 0.38<br>p<sub>adj</sub> = 0.45 |
|  | +5-9 d, saline vs clozapine (1 mg/kg) | Post-hoc pairwise, FDR adjusted | 10, 10 | Mice | p < 0.038<br>p<sub>adj</sub> = 0.075 |
|  | +5-9 d, saline vs clozapine (10 mg/kg) | Post-hoc pairwise, FDR adjusted | 10, 10 | Mice | p < 0.001<br>p<sub>adj</sub> < 0.0015 |
|  | +5-9 d, clozapine (1 mg/kg) vs clozapine (10 mg/kg) | Post-hoc pairwise, FDR adjusted | 10, 10 | Mice | p = 0.086<br>p<sub>adj</sub> = 0.15 |
| **1E** | Syllable frequency in saline different from clozapine injected (1 mg/kg) |  |  |  |  |
|  | -24 h | Permutation test | 10, 10 | Mice | p = 0.61 |
|  | +1 h | Permutation test | 10, 10 | Mice | p < 0.021 |
|  | +24 h | Permutation test | 10, 10 | Mice | p = 0.43 |
|  | +5-9 d | Permutation test | 10, 10 | Mice | p = 0.14 |
| **1F** | Syllable frequency in saline different from clozapine injected (10 mg/kg) |  |  |  |  |
|  | -24 h | Permutation test | 10, 10 | Mice | p = 0.72 |
|  | +1 h | Permutation test | 10, 10 | Mice | p < 0.001 |
|  | +24 h | Permutation test | 10, 10 | Mice | p = 0.065 |
|  | +5-9 d | Permutation test | 10, 10 | Mice | p < 0.0013 |
| **1G** | Transition matrix in saline different from clozapine injected (1 mg/kg) |  |  |  |  |
|  | -24 h | Permutation test | 10, 10 | Mice | p = 0.89 |
|  | +1 h | Permutation test | 10, 10 | Mice | p < 0.024 |
|  | +24 h | Permutation test | 10, 10 | Mice | p = 0.54 |
|  | +5-9 d | Permutation test | 10, 10 | Mice | p = 0.44 |
| **1H** | Transition matrix in saline different from clozapine injected (10 mg/kg) |  |  |  |  |
|  | -24 h | Permutation test | 10, 10 | Mice | p = 0.93 |
|  | +1 h | Permutation test | 10, 10 | Mice | p < 0.001 |
|  | +24 h | Permutation test | 10, 10 | Mice | p < 0.025 |
|  | +5-9 d | Permutation test | 10, 10 | Mice | p < 0.01 |
| **2C** | Change in correlation different from 0 |  |  |  |  |
|  | Naive | Hierarchical bootstrap | 2659401 (8) | ROI pairs (mice) | p = 0.96 |
|  | Injection day | Hierarchical bootstrap | 2665099 (8) | ROI pairs (mice) | p < 0.043 |
|  | +24 h | Hierarchical bootstrap | 2652132 (8) | ROI pairs (mice) | p = 0.54 |
|  | +48 h | Hierarchical bootstrap | 1976602 (6) | ROI pairs (mice) | p = 0.15 |
|  | +4-9 d | Hierarchical bootstrap | 2665099 (8) | ROI pairs (mice) | p < 0.001 |
|  | Change in correlation different between saline and clozapine | Hierarchical bootstrap | 2665099 (8)<br>4306608 (13) | ROI pairs (mice) | p < 0.001 |
| **3A** | Developmentally (Dev) labeled neurons depth different from adult labeled neurons | Hierarchical bootstrap | 5100 (3), 6009 (3) | Neurons (mice) | p < 0.001 |
|  | Dev labeled neurons depth different from adult labeled neurons | Paired t-test | 3, 3 | Mice | p < 0.0014 |
| **3B** | Dev labeled neurons depth different from adult labeled neurons | Hierarchical bootstrap | 7168 (4), 9214 (4) | Neurons (mice) | p < 0.001 |
|  | Dev labeled neurons depth different from adult labeled neurons | Paired t-test | 4, 4 | Mice | p < 0.001 |
| **3C** | Dev labeled neurons depth different from adult labeled neurons | Hierarchical bootstrap | 12628 (3), 7959 (3) | Neurons (mice) | p < 0.001 |
|  | Dev labeled neurons depth different from adult labeled neurons | Paired t-test | 3, 3 | Mice | p < 0.015 |
| **3G** | Change in correlation in all Tlx3 neurons different from adult labeled ones | Hierarchical bootstrap | 2316393 (7), 1670765 (5) | ROI pairs (mice) | p < 0.001 |
|  | Change in correlation in adult labeled at +6-9 d different from 0 | Hierarchical bootstrap | 1670765 (5) | ROI pairs (mice) | p < 0.013 |
| **4B** | Resting potential different between dev and adult labeled Tlx3 neurons | Hierarchical bootstrap | 12 (6), 10 (3) | Cells (mice) | p = 0.66 |
| **4D** | Input resistance different between dev and adult labeled Tlx3 neurons | Hierarchical bootstrap | 12 (6), 10 (3) | Cells (mice) | p < 0.05 |
| **4F** | Time constant different between dev and adult labeled Tlx3 neurons | Hierarchical bootstrap | 11 (6), 10 (3) | Cells (mice) | p = 0.0050 |
| **4H** | Threshold current different between dev and adult labeled Tlx3 neurons | Hierarchical bootstrap | 12 (6), 10 (3) | Cells (mice) | p = 0.010 |
| **4I** | F-I curve different between dev and adult labeled Tlx3 neurons | Linear mixed-effects model | 10,10 | Cells | Cell type:<br>p < 0.03<br>Current:<br>p < 0.0069<br>CellType*Current interaction:<br>p = 0.13 |
| **4M** | Proportion of cell types between dev and adult (labeling) Tlx3 neurons |  |  |  |  |
|  | L4/5/6 IT Car3 proportion dev vs adult | Paired t-test, FDR adjusted | 5,5 | Mice | p < 0.001<br>p<sub>adj</sub> < 0.001 |
|  | PT proportion dev vs adult | Paired t-test, FDR adjusted | 5,5 | Mice | p < 0.0035<br>p<sub>adj</sub> < 0.01 |
|  | NP/CT/L6b proportion dev vs adult | Paired t-test, FDR adjusted | 5,5 | Mice | p < 0.0015<br>p<sub>adj</sub> < 0.0056 |
|  | MGE proportion dev vs adult | Paired t-test, FDR adjusted | 5,5 | Mice | p = 0.13<br>p<sub>adj</sub> = 0.21 |
|  | L2/3 IT proportion dev vs adult | Paired t-test, FDR adjusted | 5,5 | Mice | p = 0.49<br>p<sub>adj</sub> = 0.49 |
|  | CGE proportion dev vs adult | Paired t-test, FDR adjusted | 5,5 | Mice | p = 0.26<br>p<sub>adj</sub> = 0.32 |
|  | DG/SUB/CA proportion dev vs adult | Paired t-test, FDR adjusted | 5,5 | Mice | p = 0.28<br>p<sub>adj</sub> = 0.32 |
|  | Other proportion dev vs adult | Paired t-test, FDR adjusted | 5,5 | Mice | p < 0.042<br>p<sub>adj</sub> = 0.082 |
| **5B** | Response different between saline and clozapine injected mice |  |  |  |  |
|  | -24 h | Hierarchical bootstrap | 3633 (23), 6967 (41) | Neurons (sites) | In figure for each time bin |
|  | +1 h | Hierarchical bootstrap | 3633 (23), 6967 (41) | Neurons (sites) | In figure for each time bin |
|  | +24 h | Hierarchical bootstrap | 3633 (23), 6967 (41) | Neurons (sites) | In figure for each time bin |
|  | +5-9 d | Hierarchical bootstrap | 3633 (23), 6967 (41) | Neurons (sites) | In figure for each time bin |
| **5C** | Reliability different between saline and clozapine injected mice |  |  |  |  |
|  | -24 h | Hierarchical bootstrap | 3941 (25), 6967 (41) | Neurons (sites) | p = 0.093 |
|  | +1 h | Hierarchical bootstrap | 3941 (25), 6967 (41) | Neurons (sites) | p < 0.001 |
|  | +24 h | Hierarchical bootstrap | 3633 (25), 6967 (41) | Neurons (sites) | p = 0.44 |
|  | +5-9 d | Hierarchical bootstrap | 3941 (25), 6967 (41) | Neurons (sites) | p < 0.013 |
|  | Reliability of clozapine injected mice at 5-9 d larger than at -24 h | Hierarchical bootstrap | 6967 (41) | Neurons (sites) | p < 0.001 |
| **5E** | Response amplitude different between saline and clozapine injected mice |  |  |  |  |
|  | -24 h | Hierarchical bootstrap | 136 (18), 366 (27) | Neurons (mice) | p = 0.99 |
|  | +1 h | Hierarchical bootstrap | 87 (11), 1678 (34) | Neurons (mice) | p = 0.27 |
|  | +24 h | Hierarchical bootstrap | 69 (10), 240 (26) | Neurons (mice) | p = 0.92 |
|  | +5-9 d | Hierarchical bootstrap | 114 (14), 586 (33) | Neurons (mice) | p < 0.05 |
| **5F** | Fraction of positively responding neurons different between saline and clozapine injected mice |  |  |  |  |
|  | -24 h | Hierarchical bootstrap | 136 (18), 366 (27) | Neurons (sites) | p = 0.82 |
|  | +1 h | Hierarchical bootstrap | 87 (11), 1678 (34) | Neurons (sites) | p < 0.011 |
|  | +24 h | Hierarchical bootstrap | 69 (10), 240 (26) | Neurons (sites) | p = 0.49 |
|  | +5-9 d | Hierarchical bootstrap | 114 (14), 586 (33) | Neurons (sites) | p < 0.045 |
| **6C** | Response different between saline and clozapine injected mice |  |  |  |  |
|  | -24 h | Hierarchical bootstrap | 850 (5), 1876 (10) | Neurons (sites) | In figure for each time bin |
|  | +1 h | Hierarchical bootstrap | 850 (5), 1876 (10) | Neurons (sites) | In figure for each time bin |
|  | +24 h | Hierarchical bootstrap | 850 (5), 1876 (10) | Neurons (sites) | In figure for each time bin |
|  | +5-9 d | Hierarchical bootstrap | 850 (5), 1876 (10) | Neurons (sites) | In figure for each time bin |
| **6D** | Reliability different between saline and clozapine injected mice |  |  |  |  |
|  | -24 h | Hierarchical bootstrap | 850 (5), 1876 (10) | Neurons (sites) | p = 0.19 |
|  | +1 h | Hierarchical bootstrap | 850 (5), 1876 (10) | Neurons (sites) | p = 0.55 |
|  | +24 h | Hierarchical bootstrap | 850 (5), 1876 (10) | Neurons (sites) | p = 0.61 |
|  | +5-9 d | Hierarchical bootstrap | 850 (5), 1876 (10) | Neurons (sites) | p = 0.27 |
| **6E** | Response different between saline and clozapine injected mice |  |  |  |  |
|  | -24 h | Hierarchical bootstrap | 1783 (10), 902 (6) | Neurons (sites) | In figure for each time bin |
|  | +1 h | Hierarchical bootstrap | 1783 (10), 902 (6) | Neurons (sites) | In figure for each time bin |
|  | +24 h | Hierarchical bootstrap | 1783 (10), 902 (6) | Neurons (sites) | In figure for each time bin |
|  | +5-9 d | Hierarchical bootstrap | 1783 (10), 902 (6) | Neurons (sites) | In figure for each time bin |
| **6F** | Reliability different between saline and clozapine injected mice |  |  |  |  |
|  | -24 h | Hierarchical bootstrap | 1783 (10), 902 (6) | Neurons (sites) | p = 0.27 |
|  | +1 h | Hierarchical bootstrap | 1783 (10), 902 (6) | Neurons (sites) | p = 0.28 |
|  | +24 h | Hierarchical bootstrap | 1783 (10), 902 (6) | Neurons (sites) | p = 0.42 |
|  | +5-9 d | Hierarchical bootstrap | 1783 (10), 902 (6) | Neurons (sites) | p < 0.0077 |
|  | Reliability of clozapine injected mice at 5-9 d larger than at -24 h | Hierarchical bootstrap | 1783 (10) | Neurons (sites) | p < 0.001 |
| **6H** | Response amplitude different between saline and clozapine injected mice |  |  |  |  |
|  | -24 h | Hierarchical bootstrap | 235 (10), 123 (6) | Neurons (sites) | p = 0.57 |
|  | +1 h | Hierarchical bootstrap | 195 (10), 69 (6) | Neurons (sites) | p = 0.51 |
|  | +24 h | Hierarchical bootstrap | 237 (10), 130 (6) | Neurons (sites) | p = 0.21 |
|  | +5-9 d | Hierarchical bootstrap | 342 (10), 430 (6) | Neurons (sites) | p = 0.73 |
| **6I** | Fraction of positively responding neurons |  |  |  |  |
|  | -24 h | Hierarchical bootstrap | 235 (10), 123 (6) | Neurons (sites) | p = 0.31 |
|  | +1 h | Hierarchical bootstrap | 195 (10), 69 (6) | Neurons (sites) | p = 0.40 |
|  | +24 h | Hierarchical bootstrap | 237 (10), 130 (6) | Neurons (sites) | p = 0.14 |
|  | +5-9 d | Hierarchical bootstrap | 342 (10), 430 (6) | Neurons (sites) | p = 0.20 |
| **7D** | Response different between saline and clozapine injected mice |  |  |  |  |
|  | -24 h | Hierarchical bootstrap | 7121 (44), 8552 (50) | Neurons (sites) | In figure for each time bin |
|  | +1 h | Hierarchical bootstrap | 7121 (44), 8552 (50) | Neurons (sites) | In figure for each time bin |
|  | +24 h | Hierarchical bootstrap | 7121 (44), 8552 (50) | Neurons (sites) | In figure for each time bin |
|  | +5-9 d | Hierarchical bootstrap | 7121 (44), 8552 (50) | Neurons (sites) | In figure for each time bin |
| **7E** | Reliability different between saline and clozapine injected mice |  |  |  |  |
|  | -24 h | Hierarchical bootstrap | 7429 (46), 11233 (65) | Neurons (sites) | p = 0.45 |
|  | +1 h | Hierarchical bootstrap | 7429 (46), 8693 (51) | Neurons (sites) | p < 0.024 |
|  | +24 h | Hierarchical bootstrap | 7121 (44), 8552 (50) | Neurons (sites) | p = 0.62 |
|  | +5-9 d | Hierarchical bootstrap | 7429 (44), 8693 (50) | Neurons (sites) | p = 0.49 |
| **7G** | Tuning width difference between saline and clozapine injected mice |  |  |  |  |
|  | -24 h | Hierarchical bootstrap | 1561 (45), 2004 (63) | Neurons (sites) | p = 0.53 |
|  | +1 h | Hierarchical bootstrap | 1561 (45), 1690 (49) | Neurons (sites) | p < 0.0011 |
|  | +24 h | Hierarchical bootstrap | 1466 (42), 1703 (50) | Neurons (sites) | p = 0.37 |
|  | +5-9 d | Hierarchical bootstrap | 1561 (45), 1703 (50) | Neurons (sites) | p = 0.93 |
| **8B** | Change in correlation different from 0 in aripiprazole injected mice |  |  |  |  |
|  | Naive | Hierarchical bootstrap | 2004918 (6) | ROI pairs (mice) | p = 0.95 |
|  | Injection day | Hierarchical bootstrap | 2004918 (6) | ROI pairs (mice) | p = 0.064 |
|  | +24 h | Hierarchical bootstrap | 1670765 (5) | ROI pairs (mice) | p < 0.001 |
|  | +48 h | Hierarchical bootstrap | 1670765 (5) | ROI pairs (mice) | p < 0.001 |
|  | +4-9 d | Hierarchical bootstrap | 2004918 (6) | ROI pairs (mice) | p < 0.001 |
|  | Change in correlation different between saline and aripiprazole | Hierarchical bootstrap | 2004918 (6)<br>4306608 (13) | ROI pairs (mice) | p < 0.001 |
| **8C** | Change in correlation different from 0 in risperidone injected mice |  |  |  |  |
|  | Naive | Hierarchical bootstrap | 1336612 (4) | ROI pairs (mice) | p = 0.99 |
|  | Injection day | Hierarchical bootstrap | 1336612 (4) | ROI pairs (mice) | p = 0.16 |
|  | +24 h | Hierarchical bootstrap | 1336612 (4) | ROI pairs (mice) | p = 0.49 |
|  | +48 h | Hierarchical bootstrap | 1336612 (4) | ROI pairs (mice) | p = 0.71 |
|  | +4-9 d | Hierarchical bootstrap | 1336612 (4) | ROI pairs (mice) | p < 0.001 |
|  | Change in correlation different between saline and risperidone | Hierarchical bootstrap | 1336612 (4)<br>4306608 (13) | ROI pairs (mice) | p < 0.001 |
| **8D** | Change in correlation different from 0 in haloperidol injected mice |  |  |  |  |
|  | Naive | Hierarchical bootstrap | 2337438 (7) | ROI pairs (mice) | p = 0.95 |
|  | Injection day | Hierarchical bootstrap | 2339071 (7) | ROI pairs (mice) | p = 0.57 |
|  | +24 h | Hierarchical bootstrap | 1999220 (6) | ROI pairs (mice) | p = 0.93 |
|  | +48 h | Hierarchical bootstrap | 2004918 (6) | ROI pairs (mice) | p = 0.75 |
|  | +4-9 d | Hierarchical bootstrap | 2339071 (7) | ROI pairs (mice) | p < 0.039 |
|  | Change in correlation different between saline and haloperidol | Hierarchical bootstrap | 2339071 (7)<br>4306608 (13) | ROI pairs (mice) | p < 0.013 |
| **S1B** | Fraction of time in the center different between groups. (+1 h excluded) | Two-way repeated measures ANOVA | 10,10,10 | Mice | Treatment: p = 0.84<br>Time: p = 0.42<br>Time*treatment:<br>p = 0.68 |
|  | Fraction of time in the center different between groups. | Linear mixed-effects model | 10,10,10 | Mice | Treatment: p = 0.95<br>Time: p = 0.38 |
| **S1C** | Fraction of time in the center different between groups. (+1 h excluded) | Two-way repeated measures ANOVA | 10,10,10 | Mice | Treatment: p = 0.89<br>Time: p = 0.0069<br>Time*treatment:<br>p = 0.93 |
|  | -24 h vs +24 h; All groups | Post-hoc pairwise, FDR adjusted | 10, 10 | Mice | p = 0.70<br>p<sub>adj</sub> = 0.70 |
|  | -24 h vs +5-9 d; All groups | Post-hoc pairwise, FDR adjusted | 10, 10 | Mice | p < 0.024<br>padj < 0.035 |
|  | +24 h vs +5-9 d; All groups | Post-hoc pairwise, FDR adjusted | 10, 10 | Mice | p < 0.0024<br>p<sub>adj</sub> < 0.0071 |
|  | Fraction of time in the center different between groups. | Linear mixed-effects model | 10,10,10 | Mice | Treatment: p = 0.91<br>Time: p < 0.0025 |
| **S1D** | Fraction of time in the center different between groups. (+1 h excluded) | Two-way repeated measures ANOVA | 10,10,10 | Mice | Treatment: p = 0.32<br>Time: p < 0.001<br>Time*treatment:<br>p = 0.54 |
|  | -24 h vs +24 h; All groups | Post-hoc pairwise, FDR adjusted | 10, 10 | Mice | p < 0.020<br>p<sub>adj</sub> < 0.020 |
|  | -24 h vs +5-9 d; All groups | Post-hoc pairwise, FDR adjusted | 10, 10 | Mice | p < 0.001<br>p<sub>adj</sub> < 0.001 |
|  | +24 h vs +5-9 d; All groups | Post-hoc pairwise, FDR adjusted | 10, 10 | Mice | p < 0.0015<br>p<sub>adj</sub> < 0.0022 |
|  | Fraction of time in the center different between groups. | Linear mixed-effects model | 10,10,10 | Mice | Treatment: p = 0.40<br>Time: p < 0.001 |
| **S2B** | Dev labeled neurons depth different from adult labeled Tlx3 neurons | Hierarchical bootstrap | 24896 (10), 23182 (10) | Neurons (mice) | p < 0.001 |
|  | Dev labeled neurons depth different from adult labeled Tlx3 neurons | Signed rank test | 10,10 | Mice | p < 0.0020 |
| **S3A** | Capacitance (voltage-clamp) different between dev and adult labeled Tlx3 neurons | Hierarchical bootstrap | 12 (6), 10 (3) | Neurons (mice) | p = 0.22 |
| **S3B** | Action potential amplitude different between dev and adult labeled Tlx3 neurons | Hierarchical bootstrap | 12 (6), 10 (3) | Neurons (mice) | p = 0.69 |
| **S3C** | Action potential width different between dev and adult labeled Tlx3 neurons | Hierarchical bootstrap | 12 (6), 10 (3) | Neurons (mice) | p = 0.18 |
| **S3D** | Interspike interval adaptation ratio different between dev and adult labeled Tlx3 neurons | Hierarchical bootstrap | 12 (6), 10 (3) | Neurons (mice) | p = 0.80 |
| **S3E** | AP frequency adaptation ratio different between dev and adult labeled Tlx3 neurons | Hierarchical bootstrap | 12 (6), 10 (3) | Neurons (mice) | p = 0.94 |
| **S3F** | Coefficient of variation for interspike interval different between dev and adult labeled Tlx3 neurons | Hierarchical bootstrap | 12 (6), 10 (3) | Neurons (mice) | p = 0.55 |
| **S6A** | Response amplitude greater in clozapine vs saline injected mice (reliability threshold = 0.75) |  |  |  |  |
|  | -24 h | Hierarchical bootstrap | 284 (19), 726 (32) | Neurons (sites) | In figure for each time bin |
|  | +1 h | Hierarchical bootstrap | 180 (16), 2271 (37) | Neurons (sites) | In figure for each time bin |
|  | +24 h | Hierarchical bootstrap | 182 (16), 518 (33) | Neurons (sites) | In figure for each time bin |
|  | +5-9 d | Hierarchical bootstrap | 262 (20), 1183 (34) | Neurons (sites) | In figure for each time bin |
| **S6B** | Response amplitude in saline injected greater than in clozapine injected mice | Hierarchical bootstrap | 262 (20), 1183 (34) | Neurons (sites) | p = 0.067 |
| **S6C** | Fraction in saline injected greater than in clozapine injected mice | Hierarchical bootstrap | 262 (20), 1183 (34) | Neurons (sites) | p = 0.084 |
| **S6D** | Response amplitude greater in clozapine vs saline injected (reliability threshold = 0.80) mice |  |  |  |  |
|  | -24 h | Hierarchical bootstrap | 203 (19), 548 (31) | Neurons (sites) | In figure for each time bin |
|  | +1 h | Hierarchical bootstrap | 137 (13), 2019 (37) | Neurons (sites) | In figure for each time bin |
|  | +24 h | Hierarchical bootstrap | 125 (15), 378 (30) | Neurons (sites) | In figure for each time bin |
|  | +5-9 d | Hierarchical bootstrap | 191 (20), 843 (34) | Neurons (sites) | In figure for each time bin |
| **S6E** | Response amplitude in saline injected greater than in clozapine injected mice | Hierarchical bootstrap | 191 (20), 843 (34) | Neurons (sites) | p < 0.039 |
| **S6F** | Fraction in saline injected greater than in clozapine injected mice | Hierarchical bootstrap | 191 (20), 843 (34) | Neurons (sites) | p < 0.040 |
| **S6G** | Response amplitude greater in clozapine vs saline injected (reliability threshold = 0.85) mice |  |  |  | In figure for each time bin |
|  | -24 h | Hierarchical bootstrap | 136 (18), 366 (27) | Neurons (sites) | In figure for each time bin |
|  | +1 h | Hierarchical bootstrap | 87 (11), 1678 (34) | Neurons (sites) | In figure for each time bin |
|  | +24 h | Hierarchical bootstrap | 69 (10), 240 (26) | Neurons (sites) | In figure for each time bin |
|  | +5-9 d | Hierarchical bootstrap | 114 (14), 586 (33) | Neurons (sites) | In figure for each time bin |
| **S6H** | Response amplitude in saline injected greater than in clozapine injected mice | Hierarchical bootstrap | 114 (14), 586 (33) | Neurons (sites) | p < 0.024 |
| **S6I** | Fraction in saline injected greater than in clozapine injected mice | Hierarchical bootstrap | 114 (14), 586 (33) | Neurons (sites) | p < 0.020 |
| **S6J** | Response amplitude greater in clozapine vs saline injected (reliability threshold = 0.90) mice |  |  |  |  |
|  | -24 h | Hierarchical bootstrap | 67 (16), 210 (25) | Neurons (sites) | In figure for each time bin |
|  | +1 h | Hierarchical bootstrap | 33 (7), 1233 (32) | Neurons (sites) | In figure for each time bin |
|  | +24 h | Hierarchical bootstrap | 30 (6), 134 (24) | Neurons (sites) | In figure for each time bin |
|  | +5-9 d | Hierarchical bootstrap | 69 (10), 323 (24) | Neurons (sites) | In figure for each time bin |
| **S6K** | Response amplitude in saline injected greater than in clozapine injected mice | Hierarchical bootstrap | 69 (10), 323 (24) | Neurons (sites) | p = 0.076 |
| **S6L** | Fraction in saline injected greater than in clozapine injected mice | Hierarchical bootstrap | 69 (10), 323 (24) | Neurons (sites) | p < 0.014 |
| **S6M** | Response amplitude greater in clozapine vs saline injected (-0.5 ≤ reliability ≤ 0.5) |  |  |  |  |
|  | -24 h | Hierarchical bootstrap | 3049 (25), 4980 (41) | Neurons (sites) | In figure for each time bin |
|  | +1 h | Hierarchical bootstrap | 3308 (25), 3393 (38) | Neurons (sites) | In figure for each time bin |
|  | +24 h | Hierarchical bootstrap | 2940 (23), 5285 (41) | Neurons (sites) | In figure for each time bin |
|  | +5-9 d | Hierarchical bootstrap | 3045 (25), 4143 (41) | Neurons (sites) | In figure for each time bin |
| **S6N** | Response amplitude in saline injected greater than in clozapine injected | Hierarchical bootstrap | 3045 (25), 4143 (41) | Neurons (sites) | p = 0.33 |
| **S6O** | Fraction in saline injected greater than in clozapine injected | Hierarchical bootstrap | 3045 (25), 4143 (41) | Neurons (sites) | p = 0.13 |

**Table S2.** Analyses performed with each mouse. In this table, unique mice that had the same genotype, received the same vectors with the same delivery method, and whose data contributed to the same figure, were collapsed to a single row. In cases where data from a mouse contributed to all panels in a figure, the panel indication was omitted.

| Number of mice | Genotype | AAV vectors | Delivery method | Figure |
|---|---|---|---|---|
| 30 | C57BL/6J | N/A | N/A | 1, S1 |
| 8 | Tlx3-Cre x Ai148 | N/A | N/A | 2, 3G |
| 13 | Tlx3-Cre x Ai148 | N/A | N/A | 2, 8 |
| 3 | Tlx3-Cre x Ai14 | 2/1-Ef1α-DIO-EGFP | Intracerebral injection | 3A, S2 |
| 4 | Tlx3-Cre x Ai14 | PHP.eB-Ef1α-DIO-EGFP | Intracerebral injection | 3B, S2 |
| 3 | Tlx3-Cre x Ai14 | PHP.eB-Ef1α-DIO-EGFP | Retro-orbital deposit | 3C, S2 |
| 8 | Tlx3-Cre | PHP.eB-Ef1α-DIO-GCaMP6s | Retro-orbital deposit | 3F, G |
| 6 | Tlx3-Cre x Ai14 | 2/1-Ef1α-DIO-EGFP | Intracerebral injection | 4B-I, S3 |
| 5 | Tlx3-Cre x Ai14 | 2/1-Ef1α-DIO-EGFP | Intracerebral injection | 4K-N, S4 |
| 16 | C57BL/6J | 2/1-hSyn-ChrimsonR-tdTomato<br>2/9-Ef1α-GCaMP6f | Intracerebral injection | 5, S5, S6, 7, S8 |
| 20 | C57BL/6J | 2/1-hSyn-ChrimsonR-tdTomato<br>2/9-Ef1α-GCaMP8s | Intracerebral injection | 5, S5, S6, 7, S8 |
| 8 | C57BL/6J | 2/1-hSyn-ChrimsonR-tdTomato<br>2/9-Ef1α-GCaMP6f | Intracerebral injection | 6C-D, S7, 7, S8 |
| 8 | C57BL/6J | 2/1-hSyn-ChrimsonR-tdTomato<br>2/9-Ef1α-GCaMP6f | Intracerebral injection | 6E-I, S7, 7, S8 |
| 6 | C57BL/6J | 2/9-Ef1α-GCaMP6f<br>2/1-hSyn-ChrimsonR-tdTomato | Intracerebral injection | 7, S8 |
| 1 | C57BL/6J | 2/9-Ef1α-GCaMP8s<br>2/1-hSyn-ChrimsonR-tdTomato | Intracerebral injection | 7, S8 |
| 6 | Tlx3-Cre x Ai148 | N/A | N/A | 8B |
| 4 | Tlx3-Cre x Ai148 | N/A | N/A | 8C |
| 7 | Tlx3-Cre x Ai148 | N/A | N/A | 8D |

## Notes

### Competing Interest Statement

The authors have declared no competing interest.

### Summary of Updates

We added new experiments showing that the long-term decorrelation of L5 IT network activity is not specific to clozapine: a single dose of haloperidol, risperidone, or aripiprazole produced a similar effect. These experiments also include a saline-injected control group, which shows that the decorrelation is not explained by changes in correlation over time. In addition, we revised the Summary, Introduction, and Discussion to focus on the dissociation between the time course of the effects and receptor occupancy, and rephrased the text throughout for clarity.

